# H3K36 methylation and DNA-binding both promote Ioc4 recruitment and Isw1b remodeller function

**DOI:** 10.1101/2021.02.26.432832

**Authors:** Jian Li, Lena Bergmann, Andreia Rafael de Almeida, Kimberly M. Webb, Madelaine M. Gogol, Philipp Voigt, Yingfang Liu, Huanhuan Liang, Michaela M. Smolle

**Author notes:** These authors contributed equally to this manuscript (Jian Li, Lena Bergmann).

## Abstract

The Isw1b chromatin-remodelling complex is specifically recruited to gene bodies to help retain pre-existing histones during transcription by RNA polymerase II. Recruitment is dependent on H3K36 methylation and the Isw1b subunit Ioc4, which contains an N-terminal PWWP domain. Here, we present the crystal structure of the Ioc4-PWWP domain, including a detailed functional characterization of the domain on its own as well as in the context of full-length Ioc4 and the Isw1b remodeller. The Ioc4-PWWP domain preferentially binds H3K36me3-containing nucleosomes. Its ability to bind DNA is required for nucleosome binding. It is also furthered by the unique insertion motif present in Ioc4-PWWP. The ability to bind H3K36me3 and DNA promote the interaction of full-length Ioc4 with nucleosomes *in vitro* and they are necessary for its recruitment to gene bodies *in vivo*. Furthermore, a fully functional Ioc4-PWWP domain promotes efficient remodelling by Isw1b and the maintenance of ordered chromatin *in vivo*, thereby preventing the production of non-coding RNAs.

## Introduction

DNA is packaged into chromatin, a highly condensed nucleo-protein complex that serves as a substrate for all nuclear, DNA-based processes, such as transcription, replication or DNA repair. During gene transcription by RNA polymerase II (RNAPII) nucleosomes first have to be removed in front of the enzyme and then restored after RNAPII passage, so as to prevent inappropriate access to the underlying DNA sequences by the transcription machinery (1). Organisation of chromatin structure is achieved by the concerted actions of histone modifying enzymes, chromatin remodellers and histone chaperones (2).

Many different post-translational histone modifications have been identified, including lysine acetylation and methylation. PTMs can affect chromatin organisation directly, for example by altering the electrostatic surface of histones. Alternatively, they can affect chromatin structure indirectly by providing binding platforms for specific reader modules present in many chromatin-associated factors. Over 20 such reader domains have been identified (3).

Remodelling factors use the energy generated by ATP hydrolysis to affect chromatin organisation as a consequence of nucleosome sliding, eviction or assembly (4). ISWI (*I*mitation *S*witch) is one of four families of remodelling enzymes that are conserved from yeast to humans. The Iswi remodeller was initially identified in *Drosophila melanogaster* (5) and possesses two homologs in *Saccharomyces cerevisiae*, Isw1 and Isw2 (6). In turn, Isw1 is the catalytic subunit of two distinct remodelling complexes, Isw1a and Isw1b (7). Previous work has shown that the Isw1b chromatin remodeller is involved in retaining H3K36-methylated histones and thereby maintaining proper chromatin organisation over gene bodies (8,9). More recently, Isw1b was implicated in the upkeep of regular phasing of nucleosomal arrays by resolving closely packed dinucleosomes (10). Isw1b consists of two other subunits apart from Isw1, namely Ioc2 and Ioc4 (7). Recruitment of the Isw1 remodellers to different genomic regions depends on these associated subunits. In case of Isw1b, its localisation to coding sequences depends on Ioc4. In its absence pre-existing, H3K36 trimethylated histones are lost from ORFs. Deletion of *ISW1* has the same effect (8).

The only domain annotated within Ioc4 is an N-terminal PWWP domain. PWWP domains are part of the Royal superfamily and were originally identified as DNA-binding domains (11–13). Subsequent studies revealed that most PWWP-containing proteins associate with methylated histones via a conserved aromatic cage (3). Most domains examined display a preference for histone H3 trimethylated at Lys36 (H3K36me3), although some bind to H4K20me (Pdp1, HDGF2) or H3K79me3 (HDGF2) (14,15).

While the importance of full-length Ioc4 for Isw1b recruitment and function has been demonstrated previously, we wanted to characterise in more detail the importance of the Ioc4 PWWP domain (Ioc4_PWWP_) in these processes. Therefore, we solved the crystal structure of Ioc4_PWWP_ and related its structural features to Ioc4_PWWP_ functions, both *in vivo* and *in vitro*. We found that full Isw1b remodelling activity is reached in the presence of a functional Ioc4 PWWP domain. Also, recruitment of the Isw1b remodeller through its Ioc4 subunit *in vivo* depends on interactions between Ioc4_PWWP_ and H3K36 methylation as well as DNA binding. Mutants not able to recognise either methylation and/or to bind to DNA display reduced Isw1b remodelling activities, reduced chromatin association of Ioc4 *in vivo* and increased production of non-coding RNAs. This highlights the importance of native Ioc4_PWWP_ interactions for remodeller recruitment and its function in chromatin organisation.

## Results

### The PWWP domain promotes binding of Ioc4 to nucleosomes *in vitro*

Previously, we found that localisation and binding of Ioc4 to nucleosomes depend on Set2-mediated H3K36 trimethylation both *in vivo* and *in vitro* (8). Therefore, we wanted to assess the role and importance of the Ioc4 PWWP domain in this process in more detail.

First, we repeated our previous *in vitro* binding experiments using electromobility shift assays (EMSA) with some modifications. All binding assays were performed using nucleosomal core particles (NCPs) reconstituted onto 147 bp DNA fragments rather than mononucleosomes with DNA overhangs. Furthermore, we used a peptide ligation strategy to generate histone H3 containing uniformly unmethylated (me0) or trimethylated (me3) Lys36, which were subsequently used for histone octamer assembly and NCP reconstitution instead of chemically modified histone H3 (8). We also used a different Ioc4_ΔPWWP_ construct. Previously, we had based our Ioc4_ΔPWWP_ construct on the original annotation of the Ioc4 PWWP domain (7). However, this left part of the PWWP domain intact since the crystal structure reveals a much larger PWWP domain than anticipated (see below). Instead, we generated a new Ioc4_ΔPWWP_ construct with amino acids 1-178 deleted. The different experimental setups and Ioc4 constructs yield very comparable and reproducible results (Fig.1A, Smolle et al. (8)). We find that full-length Ioc4 displays a slight but reproducible preference for trimethylated H3K36-containing nucleosome core particles when compared to unmethylated NCPs (Fig. 1A-C). This preference for H3K36 trimethylated NCPs becomes more marked when we perform a competitive binding assay using differentially labelled nucleosomes (Fig. 1C). Deletion of the PWWP domain (aa1-178) results in a considerable loss of binding of Ioc4_ΔPWWP_ to NCPs. Furthermore, Ioc4_ΔPWWP_ exhibits similar affinities towards unmethylated and trimethylated H3K36-containing nucleosome core particles (Fig. 1A,B).

**Figure 1.**
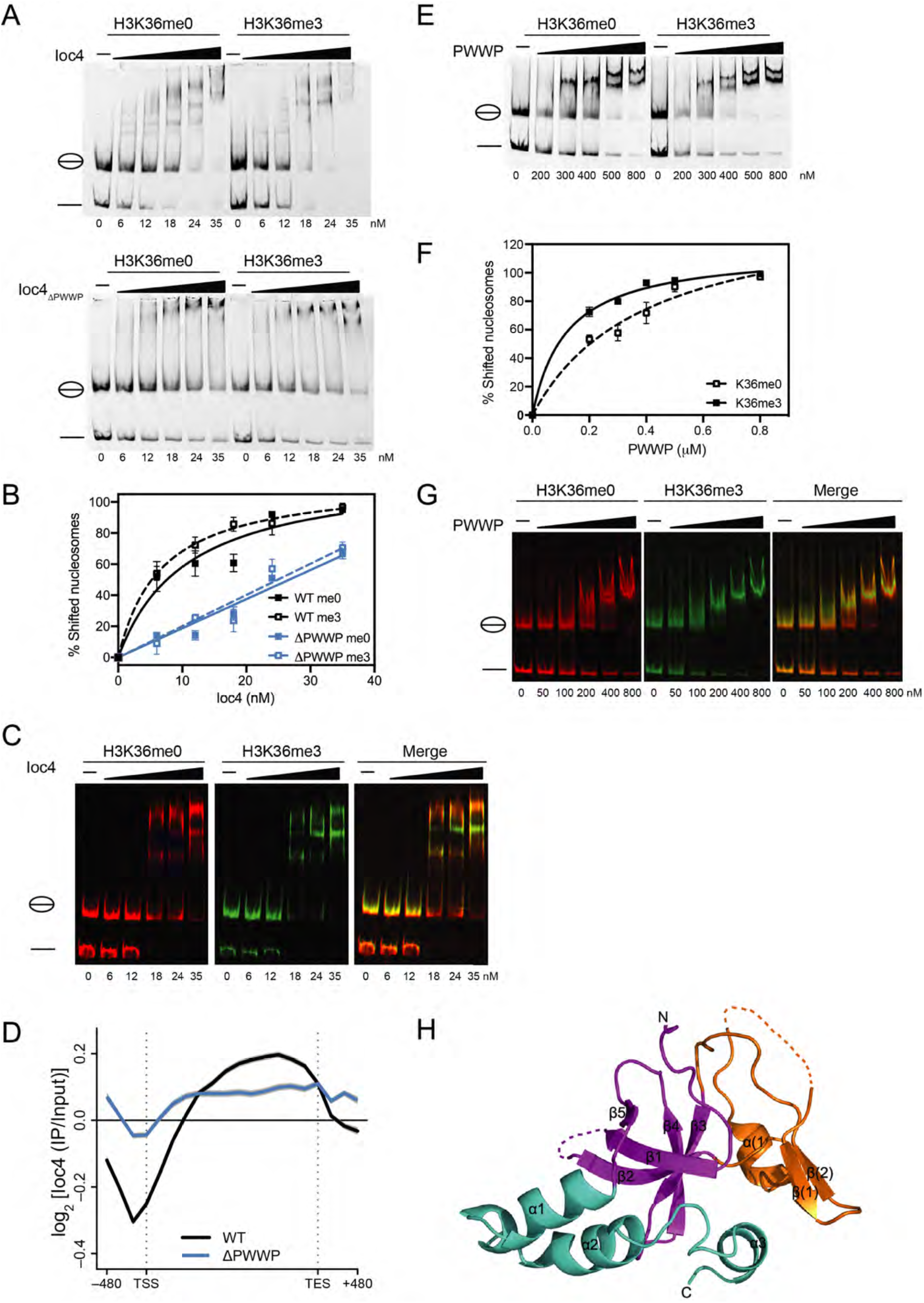
Functional Ioc4_PWWP_ is promotes chromatin binding *in vivo* and *in vitro*. **(A)** EMSA analysis of fulllength, wildtype Ioc4 and Ioc4_ΔPWWP_ binding to NCPs containing either unmethylated or trimethylated histone H3K36. **(B)** Quantitation of EMSAs shown in (A). (**C**) Competitive EMSA (cEMSA) analysis of full length, wildtype Ioc4 binding to unmethylated NCPs (red) and trimethylated (green) H3K36 NCPs. **(D)** Metagene analysis (n=6451 genes) of ChIP-chip data using yeast genome tiling arrays for wildtype (WT) Ioc4 and Ioc4_ΔPWWP_. The transcription start (TSS) and end sites (TES) are indicated. **(E)** EMSA analysis of wildtype PWWP domain binding to NCPs containing either unmethylated or trimethylated histone H3K36. **(F)** Quantitation of EMSAs shown in (E). (**G**) Competitive EMSA (cEMSA) analysis of wildtype PWWP binding to unmethylated (red) and trimethylated (green) H3K36 NCPs. **(H)** Crystal structure of the Ioc4 PWWP domain. The insertion motif between β2 and β3 is shown in orange.

### PWWP domain deletion results in reduced targeting of Isw1b *in vivo*

In wildtype cells Ioc4 localises to the mid- to 3’-regions of ORFs that are characterised by high levels of H3K36 methylation as shown previously by us and others (Fig. 1D) (8,16). In order to assess the contribution of the Ioc4 PWWP domain in correctly targeting Ioc4, and by inference Isw1b correctly *in vivo*, we determined the localisation of Ioc4 with and without its PWWP domain on a genome-wide level. Deletion of the Ioc4 PWWP domain does not interfere with protein folding as the mutant protein is still able to associate with Isw1 and Ioc2 to form Isw1b complex (Fig.S1A). Ioc4 without a functional PWWP domain (ΔPWWP) still localises to ORFs. However, it does so at significantly reduced levels and loses the preference for mid- to 3’-ORF regions (Fig. 1D). These more non-specific interactions are presumably mediated by other binding sites still present on the remaining Ioc4 molecule and/or interaction surfaces on the other complex subunits, such as the SANT and SLIDE domains of Isw1 (17). Taken together, these results clearly show that the Ioc4 PWWP domain promotes binding to nucleosomes *in vitro* as well as targeting Isw1b *in vivo*. Therefore, we set out to analyse the Ioc4 PWWP domain more closely and dissect its contributions to Isw1b function and chromatin association.

### The Ioc4 PWWP domain preferentially binds trimethylated H3K36 nucleosomes *in vitro*

The PWWP domain alone also preferentially binds to H3K36me3-containing NCPs *in vitro* (Fig. 1E-G). This preference is mediated by the aromatic cage in the PWWP domain (Fig. S2) and is abrogated by mutating the aromatic cage residue W22 to alanine. Similar observations have been made for other PWWP-containing proteins, such as human LEDGF, human BRPF1 or *S.p*. Hrp3 (18–20). Increased binding to the methylated over the unmethylated substrate is more apparent for the PWWP domain than for full-length Ioc4 (Fig. 1B,F). This is due to the fact that the affinity towards NCPs is considerably lower for the PWWP domain alone than that for full-length Ioc4, in agreement with our finding that the remaining Ioc4 protein contributes to nucleosome binding both *in vivo* and *in vitro* (Fig. 1B,D).

### The Ioc4 PWWP structure reveals highly charged patches and a unique insertion motif

In order to further our understanding of the Ioc4 PWWP domain we solved the crystal structure of the Ioc4 PWWP domain at 2.3 Å (Fig. 1H, Table S1). Analysis of the structure showed a high degree of structural conservation overall when compared to other PWWP domains. Overlay of the PWWP domains of Ioc4 and BRPF1 confirmed the high degree of conservation of PWWP domains at both, a structural (Fig. S3A) and sequence level (Fig. S3B). In common with other PWWP domains the Ioc4 PWWP domain contains a highly conserved, antiparallel, five-stranded β-barrel at its N-terminus. The a-helical composition at the C-terminus of the domain is also maintained, although the number and precise spatial relationship to the rest of the domain can vary in individual PWWP domains (3). A unique feature of the Ioc4 PWWP domain is the presence of an exceptionally long insertion motif (aa 27-110). Other PWWP domain-containing proteins have either no (e.g. LEDGF) or a much shorter (e.g. BRPF1) insertion motif (3,21).

Close analysis of the Ioc4 PWWP domain crystal structure revealed the presence of one acidic and two basic patches (Fig. 2A). PWWP domains were initially identified as DNA-binding domains and several PWWP domains have been shown to bind to double-stranded DNA (18,22–25). Given this charge distribution on the Ioc4 PWWP domain we hypothesised that it may be able to bind histones and DNA directly via the acidic and basic patches, respectively. Using pull-down assays we could show that the Ioc4 PWWP domain interacts both with histone octamers (Fig. 2C) and histone H3/H4 tetramers (Fig. S4A). In both cases full-length Ioc4 displays higher affinity towards histones compared to the PWWP domain alone.

**Figure 2.**
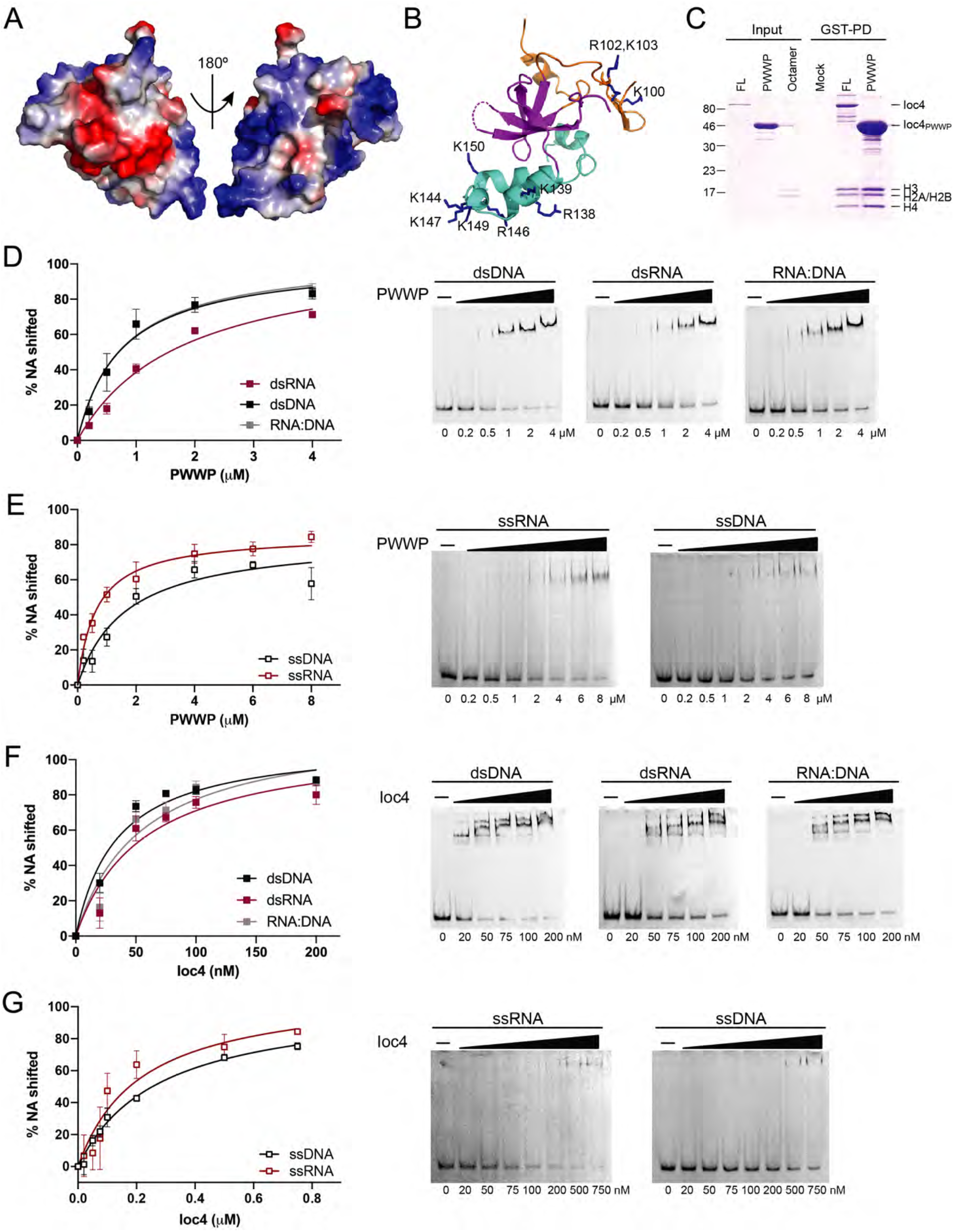
The Ioc4 PWWP domain binds double-stranded and single-stranded nucleic acids. **(A)** Electrostatic surface of the Ioc4 PWWP domain. Negative, positive and neutral charges are shown in red, blue and white, respectively. **(B)** Lysine and arginine residues contributing to the basic patches. All residues indicated were mutated and tested for DNA binding (Fig. S4D). **(C)** Pull-down assay of GST-tagged Ioc4 proteins with core histone octamers. **(D-G)** EMSA analyses of the wildtype PWWP domain (D,E) or full-length Ioc4 (F,G) binding to double stranded (D,F) or single stranded (E,G) nucleic acids.

### The Ioc4 PWWP domain binds single and double stranded nucleic acids

Next we investigated binding of the PWWP domain to DNA. Using EMSAs we could show that the Ioc4 PWWP domain can indeed bind to DNA (Fig. 2D). Since most PWWP domains do not exhibit sequence specificity (21) and are thought to interact with the negatively charged phosphate backbone we also tested binding of the domain to double-stranded (ds) RNA and RNA:DNA hybrid molecules. The PWWP domain bound both dsDNA and DNA:RNA hybrid molecules equally well, but displayed lower affinity towards dsRNA (Fig. 2D). The GC content does not seem to affect binding since both substrates are bound at equal levels (Fig. S4C). We then also tested binding of the PWWP domain to singlestranded (ss) DNA and RNA. Again it bound both molecules, but displayed higher affinity for ssRNA (Fig. 2E). Notably, the affinity of the Ioc4 PWWP domain for dsDNA is similar to that for ssRNA (Table 1). Whether this finding has functional implications for Isw1b is unclear at present. Interestingly, previous UV cross-linking experiments showed that both Isw1 and Ioc2 interact with RNA *in vivo* (26). Overall, the PWWP domain binds H3K36 trimethylated nucleosome core particles with ca. 6-fold higher affinity compared to dsDNA, suggesting that H3K36 trimethylation, histone and DNA binding are all important for the interaction of the PWWP domain with nucleosomes.

**Table 1.**
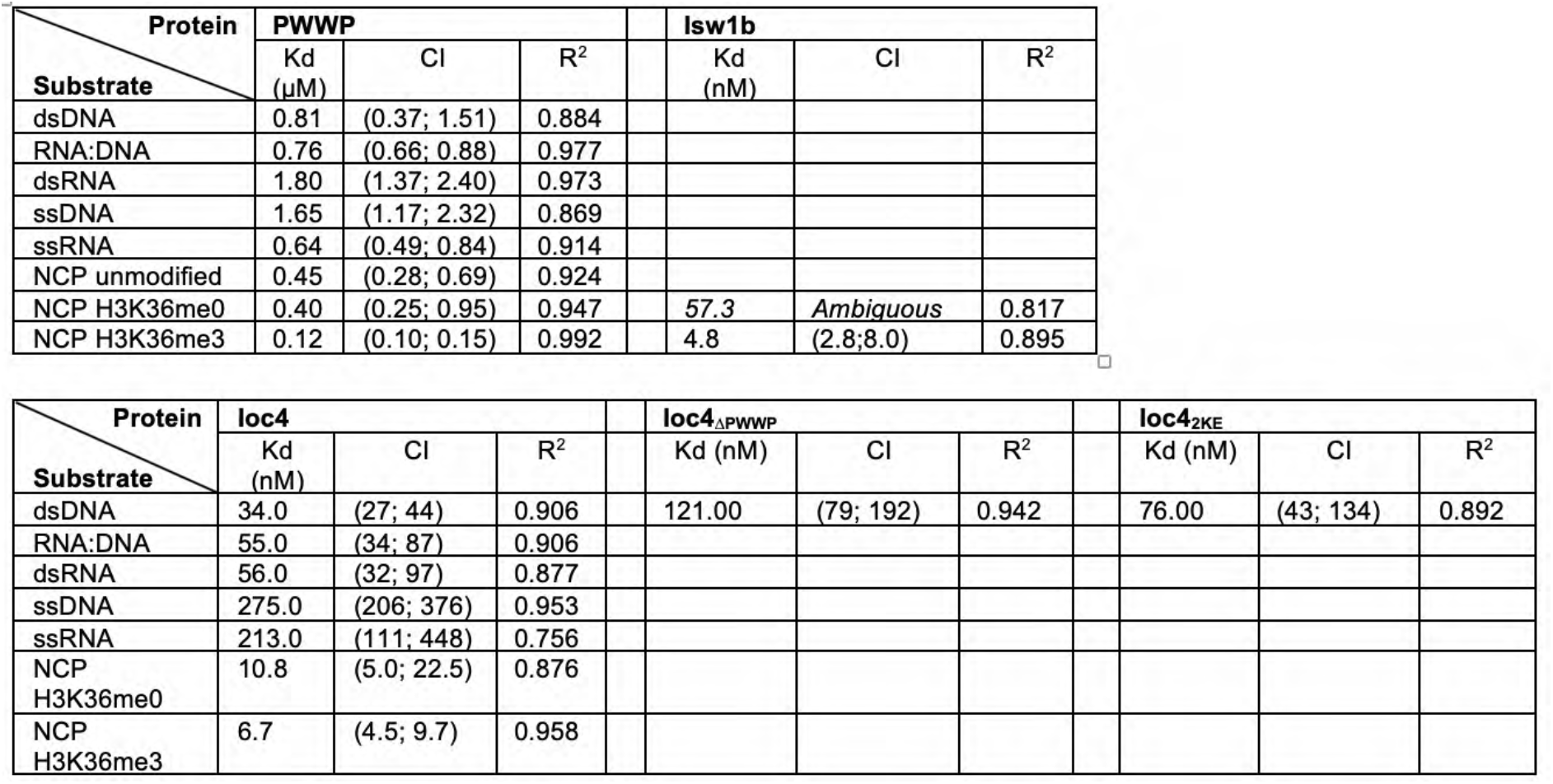
Binding parameters. The protein construct or complex is indicated. Nucleic acid substrates include single-stranded (ss) and double-stranded (ds) RNA, DNA, RNA:DNA hybrid molecules or nucleosomal core particles (NCP). Apparent dissociation constants (K_D_) are indicated if determined, including the 90% confidence interval (CI) and goodness of fit (R^2^).

We also tested interaction of full-length Ioc4 with nucleic acids. Overall, Ioc4 has a much higher affinity for all substrates when compared to the PWWP domain alone. However, the differences in affinity of Ioc4 for dsDNA and dsRNA are smaller (Fig. 2F). The same applies to Ioc4 binding to ssRNA and ssDNA (Fig. 2G).

### Nucleosome binding by Ioc4_PWWP_ requires interaction with DNA

The basic patches apparent on the Ioc4 PWWP domain consist of a series of lysine and arginine residues (Fig. 2B). We chose several surface residues to generate site mutants. We then tested the ability of the mutant proteins to bind DNA. While the wildtype PWWP protein associated with DNA, most mutants did not bind DNA *in vitro* (Fig. 3A, S4D). We then tested the binding behaviour of a PWWP domain containing K149E K150E (2KE) mutation in more detail. The 2KE mutant was completely unable to bind double-stranded DNA (Figs. 3A,B).

**Figure 3.**
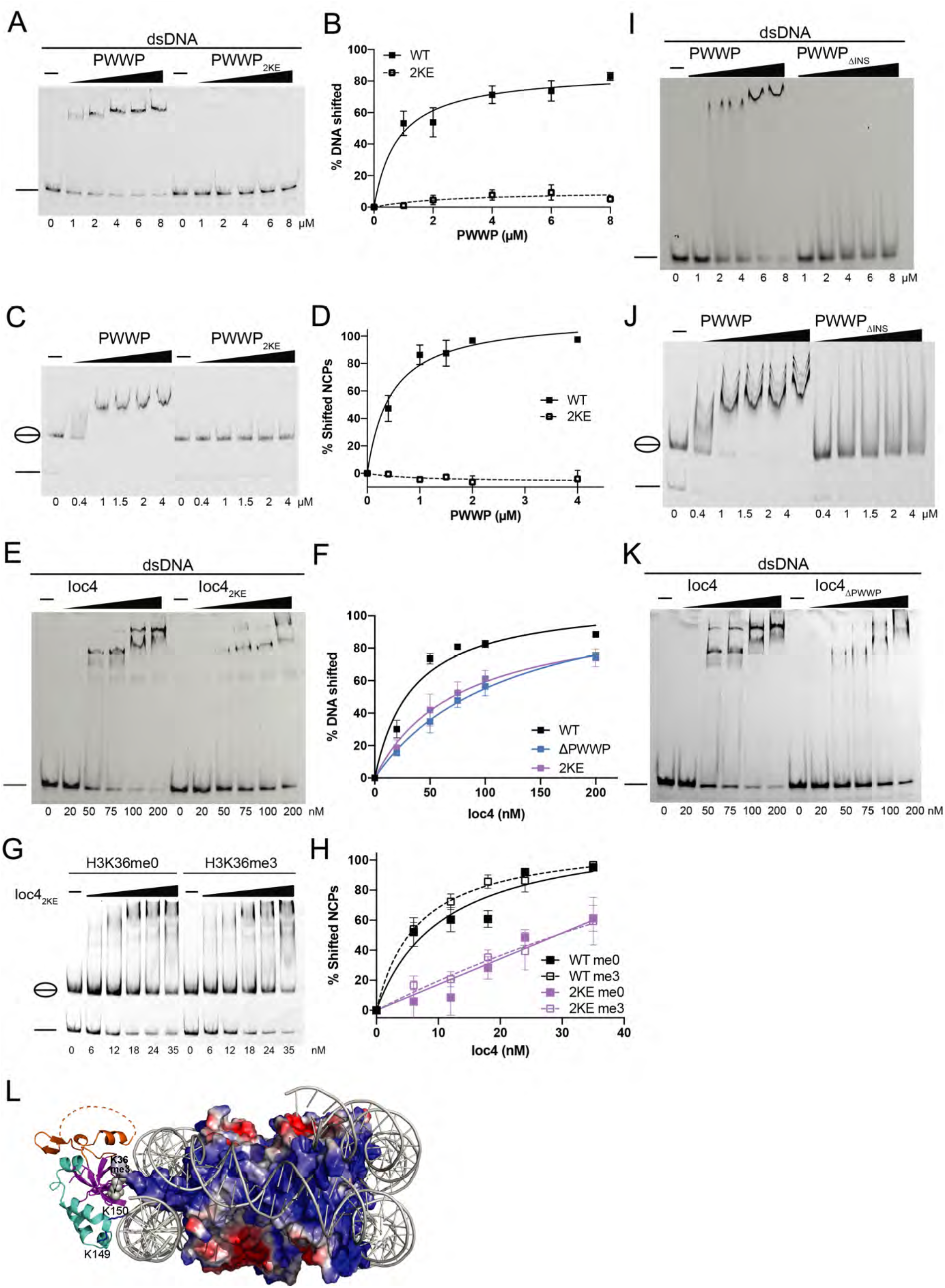
DNA-binding by Ioc4_PWWP_ promotes its interaction with nucleosomes. **(A,B,C,D,I,J)** EMSA analyses of wildtype PWWP, the PWWP_2KE_ and PWWP_δINS_ mutants. (A,C,I,J) binding to double stranded DNA (A,I) or NCPs (C,J). Quantitations of triplicate experiments are shown in (B,D). **(E,F,G,H,K)** EMSA analyses of fulllength Ioc4, the Ioc4_2KE_ and Ioc4_ΔPWWP_ mutants (E,K,G) binding to double stranded DNA (E,K) or NCPs (G). Quantitations of triplicate experiments are shown in (F,H). (**L**) Model of Ioc4 PWWP binding to NCP. Homology model based on Wang et al. (27).

Next, we tested the ability of DNA binding deficient PWWP domain to associate with nucleosomes. Surprisingly, the PWWP_2KE_ mutant exhibited no interaction with nucleosome core particles *in vitro* (Figs. 3C,D). These results suggest that DNA binding is essential in mediating the stable interaction of the Ioc4 PWWP domain with nucleosomes. Mutating K149 and K150 to alanine instead of lysine results in the same loss of binding of the mutant PWWP2KA domain to both DNA and nucleosomes (Fig. S4E,F).

### The full-length Ioc4_2KE_ mutant shows reduced binding of DNA and NCPs *in vitro*

We also tested the effects of the 2KE mutant in the context of the full-length Ioc4 protein. While the mutant displays some affinity towards DNA, the binding profile of Ioc4_2KE_ to DNA is virtually identical to that of Ioc4_ΔPWWP_ (Fig. 3E,F,K). Also, Ioc4_2KE_ does bind to nucleosome core particles, however it does so at much reduced levels (Fig. 3G,H). Furthermore, it can no longer distinguish between unmethylated and trimethylated H3K36-containing NCPs, in a manner similar to Ioc4_ΔPWWP_ (Fig. 1A,B), suggesting that the mutation of these two lysine residues is sufficient to reduce nucleosome binding of the PWWP domain even in the context of the full-length protein. The residual association of Ioc4_2KE_ with NCPs is therefore likely mediated by regions of the Ioc4 protein outside of the PWWP domain.

Two previous studies investigating the structure of the human LEDGF PWWP domain also showed the ability of this domain to bind DNA (18,19). PWWP domains in other proteins such as Pdp1, Hrp3, HDGF, MSH6, ZMYND11 and murine Dnmt3b were also shown to bind DNA (18–20,22–24). Thus, it seems the ability of PWWP domains to bind DNA is a conserved feature of PWWP domains in general. However, in contrast to our results, van Nuland and co-workers (18) found that most mutations interfering with DNA binding reduced, but did not completely abolish nucleosome binding by the LEDGF-PWWP. A further difference between the PWWP domains from Ioc4 and LEDGF is the fact that some LEDGF-PWWP DNA-binding mutants still retain their preference for H3K36-methylated nucleosomes, while the Ioc4 PWWP_2KE_ mutant does not bind to nucleosomes at all, irrespective of its H3K36 methylation status, while the full-length Ioc4_2KE_ mutant binds both types of NCPs equally well (Fig. 3G,H,S4G). Only the wildtype PWWP domain or the full-length Ioc4 protein can achieve this distinction. In part these differences may also be attributable to the fact that the authors used GST-fusion proteins for their EMSAs, which can dimerise through the GST tag and thus provide a second binding site, thereby increasing its affinity, while we used proteins containing monomeric MBP- or His-tags.

### The insertion motif promotes binding to nucleosomes, histones and DNA

Since the Ioc4 PWWP domain contains a unique insertion motif (INS) that is either completely absent or much shorter in all other PWWP domains studied so far, we sought to investigate its contributions towards Ioc4 PWWP functions. Therefore, we generated a PWWP mutant that lacks most of the insertion motif (Δ43-105, DINS). Residues for deletion were chosen to minimise potential deleterious effects on the folding of the mutant PWWP domain. Circular dichroism and/or nano-differential scanning fluorimetry (nano-DSF) showed that all purified, recombinant PWWP and Ioc4 proteins were indeed folded (Fig. S1B-D). Also, deletion of the insertion motif within the *IOC4* gene in yeast still allows for the successful purification of the mutant Isw1b_ΔINS_ remodeller, a further indication that the mutant protein is able to fold correctly and does not interfere with the binding of the Isw1 and Ioc2 subunits (Fig. S1A).

First, we tested binding of PWWP_ΔINS_ to nucleosome core particles. In contrast to the wildtype PWWP domain, deletion of the insertion motif largely abrogates stable interactions with NCPs (Fig. 3J), suggesting the insertion motif stabilises contacts between the PWWP domain and nucleosomes. Analysis of the amino acid composition of the insertion motif reveals the presence of a series of negatively charged amino acids, suggesting a potential role in histone binding. Indeed, a PWWP_ΔINS_ mutant does not bind histones at all, while the insertion motif by itself still associates with histones (Fig. S4A,B). The insertion motif also contributes several residues towards one of the basic patches seen in the crystal structure. Therefore, we also tested the ability of PWWP_ΔINS_ to bind to DNA. Unlike the wildtype domain, the PWWP_ΔINS_ mutant is not able to form a stable complex with double-stranded DNA, although it may form transient interactions at very low levels as evidenced by increased smearing of the DNA band at higher protein concentrations (Fig. 3I).

### The Ioc4 PWWP domain binds across DNA gyres

The effects of the PWWP_2KE_ and PWWP_ΔINS_ mutants can be explained by modeling our structure of the Ioc4 PWWP domain onto the recently published cryo-EM structure determined for the human LEDGF-PWWP domain bound to a nucleosome (Fig. 3L,S5) (27). We see a high degree of overlap, especially over the highly conserved beta barrel. Binding of the Ioc4 PWWP domain also occurs across DNA gyres and is mediated by the basic patch seen in Fig. 2A. Residues K149 and K150 are in close contact with the DNA sugar phosphate backbone which explains why their replacement completely abrogates DNA and nucleosome binding. Similarly, the insertion motif is in close contact with one of the nucleosomal DNA gyres, thus potentially stabilizing the interactions between the PWWP domain and the nucleosome core particle. While the stretch of negatively charged residues within the insertion motif is not resolved in the crystal structure, it seems likely that it interacts with the positively charged histone tails and thereby provide further stabilization of the interaction. A second set of positively charged residues within the insertion motif points away from the NCP surface and may contribute towards the binding of linker DNA and/or the DNA associated with neighbouring nucleosomes (Fig. S5).

### Targeting of Ioc4 *in vivo* depends on H3K36 methylation and DNA binding

The results from our *in vitro* experiments suggest that both the insertion domain and DNA binding by the Ioc4 PWWP domain promote stable interactions with DNA, histones and nucleosomes. Therefore, we wanted to test how these features influence the localisation of Ioc4 *in vivo*. To assess the effects of these mutations *in vivo*, we generated full-length, 3xFLAG-tagged Ioc4 yeast strains carrying either the K149E K150E (2KE) mutation or the insertion motif deletion (ΔINS). Furthermore, we wanted to compare the localisation of these Ioc4 mutants to the recruitment deficiencies of wildtype Ioc4 in a *set2Δ* background that contains no H3K36 methylation, as well as a yeast strain with the entire PWWP domain deleted (ΔPWWP). Finally, we assessed the impact of the PWWP aromatic cage directly by mutating W22 to alanine. Neither deletion of the PWWP domain nor the mutations of K149/K150 and/or W22 affected Ioc4 protein expression or stability when compared to the wildtype protein (Fig. S6A).

We then determined the genome-wide localisation of Ioc4 in a *set2Δ* background as well as of the Ioc4_2KE_ and Ioc4_ΔINS_ mutants, and compared these to the localisation patterns of wildtype Ioc4. In the absence of H3K36 methylation (*set2*Δ) we saw reduced targeting of Ioc4 specifically to mid- and 3’-ORFs, in a manner comparable to the deletion of the entire Ioc4 PWWP domain (Fig. 4A,B,S6B). Some non-specific background binding of Ioc4 remained in the *set2Δ* mutant. The aromatic cage mutant Ioc4w22A which is unable to distinguish un- and trimethylated H3K36 nucleosomes displayed a similar reduction in ORF localisation when tested by ChIP-qPCR (Fig. 4C,S6C). The Ioc4_2KE_ mutation also resulted in reduced targeting of Ioc4 to mid- and 3’-ORFs, in a manner almost identical to that of Ioc4 localisation in a *set2Δ* mutant and comparable to the deletion of the entire Ioc4 PWWP domain (Fig. 4A-C,S6B). The background binding seen in all mutants is consistent with the presence of other interaction surfaces remaining both within the PWWP domain as well as on the remainder of the Ioc4 protein, in agreement with our *in vitro* binding assays. Furthermore, additional binding interfaces present within Isw1b on Ioc2 and/or Isw1 could also contribute to recruitment *in vivo*. Similarly, *in vitro* binding assays of purified Isw1b remodeller show preferential binding of H3K36me3-containing nucleosomes, yet can still interact with unmethylated nucleosomes albeit with lower affinity (Fig. 4D). We also tested whether Ioc4 interaction with methylated H3K36 and simultaneous DNA binding display additive effects with regards to Ioc4 localisation by generating an Ioc4_W22A 2KE_ double mutant. Using ChIP-qPCR we see no further reduction in chromatin association of the double mutant when compared to the single mutants (Fig. 4C). Taken together, these results suggest that both recognition of H3K36 methylation via the aromatic cage as well as the ability of the Ioc4 PWWP domain to bind DNA are each necessary for the correct targeting of Ioc4 and consequently of Isw1b *in vivo*.

**Figure 4.**
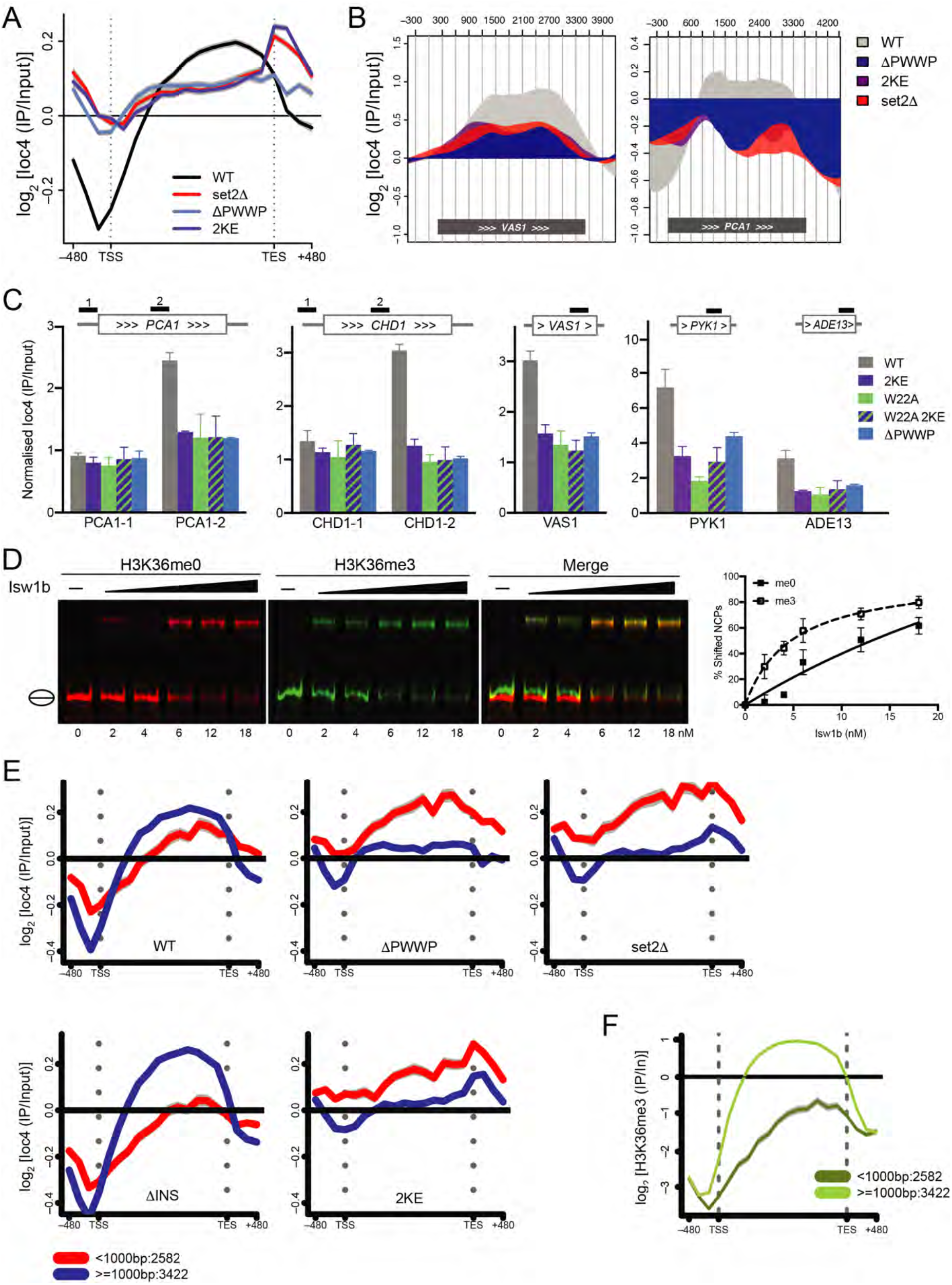
Ioc4 recruitment depends on binding to DNA and H3K36 methylation. **(A)** Metagene analysis of ChIP-chip experiments using yeast genome tiling arrays. Whole-genome average data (n=6451 genes) for three independent experiment were plotted as mean ± s.e.m. (gray) for full-length Ioc4 in wildtype (WT) and *set2Δ* backgrounds as well as Ioc4 without its PWWP domain (ΔPWWP) and the DNA-binding mutant Ioc4_2KE_. **(B)** Localisation of wildtype and mutant Ioc4 over individual genes. **(C)** ChIP-qPCR analysis of wildtype Ioc4, Ioc4_ΔPWWP_, the DNA-binding mutant Ioc4_2KE_, the aromatic cage mutant Ioc4_W22A_ and the double mutant Ioc4_W22A 2KE_. Three biological replicates were analysed and plotted as mean ± s.e.m. **(D)** cEMSA analysis of Isw1b binding to unmethylated (red) and trimethylated (green) H3K36 NCPs. **(E-F)** Metagene analysis of wildtype and mutant Ioc4 (**E**) and H3K36me3 **(F)** localization using ChIP-chip experiments and yeast genome tiling arrays. Wholegenome average data for three independent experiment were plotted as mean ± s.e.m. (gray) for genes clustered according to gene length. The number of genes in each group is indicated.

Deletion of the insertion motif alone had little overall effect on the genome-wide Ioc4 localisation over ORFs *in vivo* (Fig. S6D). Interestingly, a closer look at the data reveals that deletion of the Ioc4 insertion motif has opposite effects on Ioc4_ΔINS_ localisation for different categories of genes. While less Ioc4_ΔINS_ is found over shorter genes, increased levels of Ioc4_ΔINS_ are observed over the ORFs of genes longer than 1000 bp (Fig. 4E,S6E), suggesting multiple recruitment mechanisms may be at play for different categories of genes. Longer genes generally display more extensive domains of H3K36 trimethylation since the Set2 methyltransferase is recruited to RNAPII phosphorylated on Ser2 of its C-terminal domain (CTD). Furthermore, Set2 as well as H3K36me3 signals are mostly found over genes beyond 400 bp from the transcription start site (28,29). Indeed, looking at H3K36me3 levels in these same gene groups, we see that they are only enriched for genes longer than 1000 bp (Fig. 4F). Accordingly, loss of H3K36 trimethylation in a *set2Δ* mutant leads to reduced Ioc4 localisation precisely over this group of genes (Fig. 4E). Similarly, deletion of the Ioc4 PWWP domain leads to a similar reduction of Ioc4_ΔPWWP_ over the same set of genes (Fig. 4E). Deletion of the insertion motif leads to a significant reduction in PWWP binding to histones and nucleosomes *in vitro* (Fig. 3J, S4A). In contrast, deletion of the insertion motif in the context of full-length Ioc4 leads to a significant change in its binding behavior towards nucleosome core particles *in vitro*, resulting in the more prominent association of multiple copies of the mutant Ioc4 protein with NCPs when compared to the wildtype protein (Fig. S4I). It is possible that this is also reflected in the increased Ioc4_ΔINS_ association *in vivo* with the H3K36 trimethylated, nucleosome-packed ORFs over longer genes that are only partially disassembled during transcription. In contrast, the group of genes shorter than 1000 bp contains many highly active genes characterised by poorly organised chromatin where nucleosomes are more likely to be completely displaced from the DNA (30–33).

### Reduced targeting of Ioc4 results in growth defects and non-coding transcription

To assess the importance of functional Isw1b remodeller *in vivo* we used growth assays to investigate the impact of our mutants. Deletion of *IOC4* leads to reduced growth on media containing propiconazole, a fungicide that impairs ergosterol biosynthesis (34). Deletion of the Ioc4 PWWP domain leads to a similar growth defect as for *ioc4Δ*. Also, the aromatic cage mutant *IOC4_W22A_* and the DNA-binding deficient *IOC4_2KE_* exhibit slow growth phenotypes which are not further exacerbated in the *IOC4_W22A 2KE_* double mutant (Fig. 5A).

**Figure 5.**
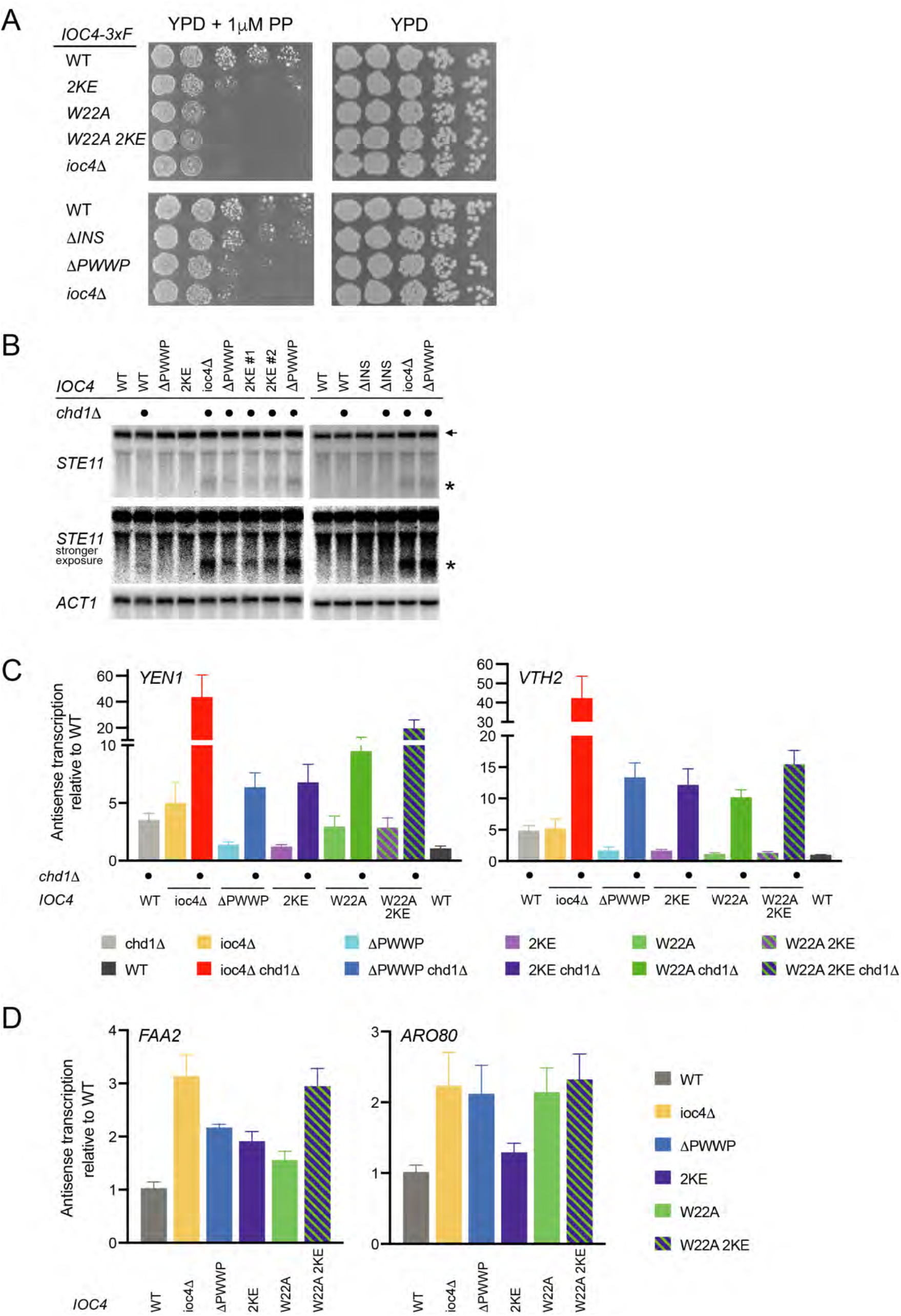
Full remodeller functions and suppression of cryptic transcription depend on Ioc4_PWWP_. **(A)** Wildtype and mutant yeast strains were grown at 30°C on YPD ± 1 μM propiconazole. **(B)** Northern blot of total RNA to investigate cryptic transcription. A probe directed against the 3’ end of *STE11* was used. *ACT1* was used as a loading control. The additional deletion of *CHD1* is indicated. The full-length (←) and short cryptic (*) transcripts are indicated. A stronger exposure better reveals the rise of cryptic transcripts in the mutants. **(C,D)** Strand-specific, multiplex RT-qPCR was used to determine the levels of antisense transcription in wildtype yeast and different *IOC4* mutants. The additional deletion of *CHD1* in (C) is indicated. Five independent experiments were performed and plotted as mean ± s.e.m.

Previous work has shown that functional Isw1b remodeller is necessary for the retention of existing histones over ORFs. Deletion of *IOC4* or *ISW1* leads to increased incorporation of soluble histones and disrupts chromatin organisation over ORFs, which in turn results in the exposure of cryptic promoters and the production of non-coding transcripts (8). Since our results suggested that the ability of the Ioc4 PWWP domain to bind nucleosomal DNA and/or to associate with methylated H3K36 is important for its recruitment, we wanted to determine whether reduced localisation of Ioc4 and Isw1b over ORFs also results in non-coding transcription. We used *STE11*, a well-characterised cryptic transcript model gene to assess the impact of our different *IOC4* mutants. Deletion of *IOC4* alone does not produce a cryptic transcript phenotype discernible by Northern blotting. However, it has been shown previously that deletion of *ISW1* and *CHD1* has additive effects with respect to cryptic transcription (8,35). Hence, we investigated the effects of various *IOC4* mutants on cryptic transcription in a *chd1Δ* background. While the overall levels of cryptic transcripts were relatively low, we observed similar and reproducible levels of the small cryptic *STE11* transcript for *IOC4*Δ, the *IOC4_2KE_* as well as the *IOC4_ΔPWWP_* mutants in a *chd1*Δ background (Fig. 5B). These results confirm that Isw1b activity in these *IOC4* mutants is reduced *in vivo*, consistent with the lower levels of mutant Ioc4 chromatin association observed. Deletion of the insertion motif leads to a minimal effect on cryptic transcription only detectable at high contrast. However, recruitment of Ioc4_ΔINS_ to *STE11* is unchanged relative to wildtype Ioc4, suggesting that increased cryptic transcription is unlikely to occur in this context.

Furthermore, we developed a new, strand-specific, multiplex RT-qPCR approach to assess the levels of antisense transcription over several genes. Candidate genes were identified from RNA-seq data obtained for *isw1*Δ. The levels of antisense (as) transcripts produced in different *IOC4* mutants were determined in either a wildtype or *chd1*Δ background (Fig. 5C,D). Chd1 has no effect on antisense transcription over the *FAA2* and *ARO80* genes. Deletion of the PWWP domain, the absence of the aromatic cage (W22A) or DNA-binding (2KE) all resulted in the production of antisense transcripts when compared to wildtype yeast (Fig. 5D). Double deletion of *IOC4* and *CHD1* shows large additive effects on antisense transcription over *YEN1* and *VTH2*. However, deletion of the PWWP domain as well as the 2KE and W22A mutants all display upregulated levels of antisense transcripts when compared to *chd1*Δ alone (Fig. 5C). The relative effects of the 2KE and W22A mutations vary depending on the asRNA investigated. With the exception of *FAA2* the effects of these two mutations are not additive, in agreement with our other *in vivo* data on growth and Ioc4 localisation.

### Ioc4 PWWP domain contributes to efficient nucleosome sliding by Isw1b

We have shown previously that cryptic transcription occurs in catalytically inactive Isw 1 mutants, indicating that remodelling activity is required to suppress this phenotype (8). The *in vivo* results in this study show that a functional Ioc4 PWWP domain is needed for correct targeting of the Isw1b remodeller and promotes suppression of non-coding transcription. We therefore wanted to test whether the Ioc4 PWWP domain is needed primarily only for the correct targeting of Isw1b remodellers to ORFs, or whether it also affects Isw1b remodelling activity directly. Therefore, we purified wildtype and mutant Isw1b remodellers from yeast cells via TAP-tagged Ioc4 subunits (Fig.S1A) and tested their remodelling activities *in vitro* using methyl-lysine analogue (MLA) H3K36Cme3-containing nucleosomes (Fig. 6). MLA nucleosomes were used for the remodeling assays because the peptide-ligated variety used for all other *in vitro* assays tended to disassemble during remodeling reactions, possibly due to the fact that peptide ligation required mutagenesis of T45 in histone H3 which is near the nucleosome dyad axis. Wildtype Isw1b preferentially moves histone octamers from a central position towards the ends of the DNA (36). Remodeller with its entire Ioc4 PWWP domain deleted (Isw1b_ΔPWWP_), as well as Isw1b containing the DNA-binding deficient Ioc4_2KE_ or the aromatic cage mutant Ioc4_W22A_ all exhibit reduced remodelling activities compared to wildtype Isw1b, although this is most apparent for the W22A mutants which displays barely any activity at all (Fig. 6A-E). It is not entirely clear why the aromatic cage mutant has a larger effect than deletion of the entire PWWP domain itself, especially since the W22A site mutation does not interfere with PWWP domain folding (Fig. S1D) or purification of the mutant remodelling complex (Fig. S1A). These results clearly show that a functional PWWP domain in Ioc4 promotes full Isw1b remodeller function.

**Figure 6.**
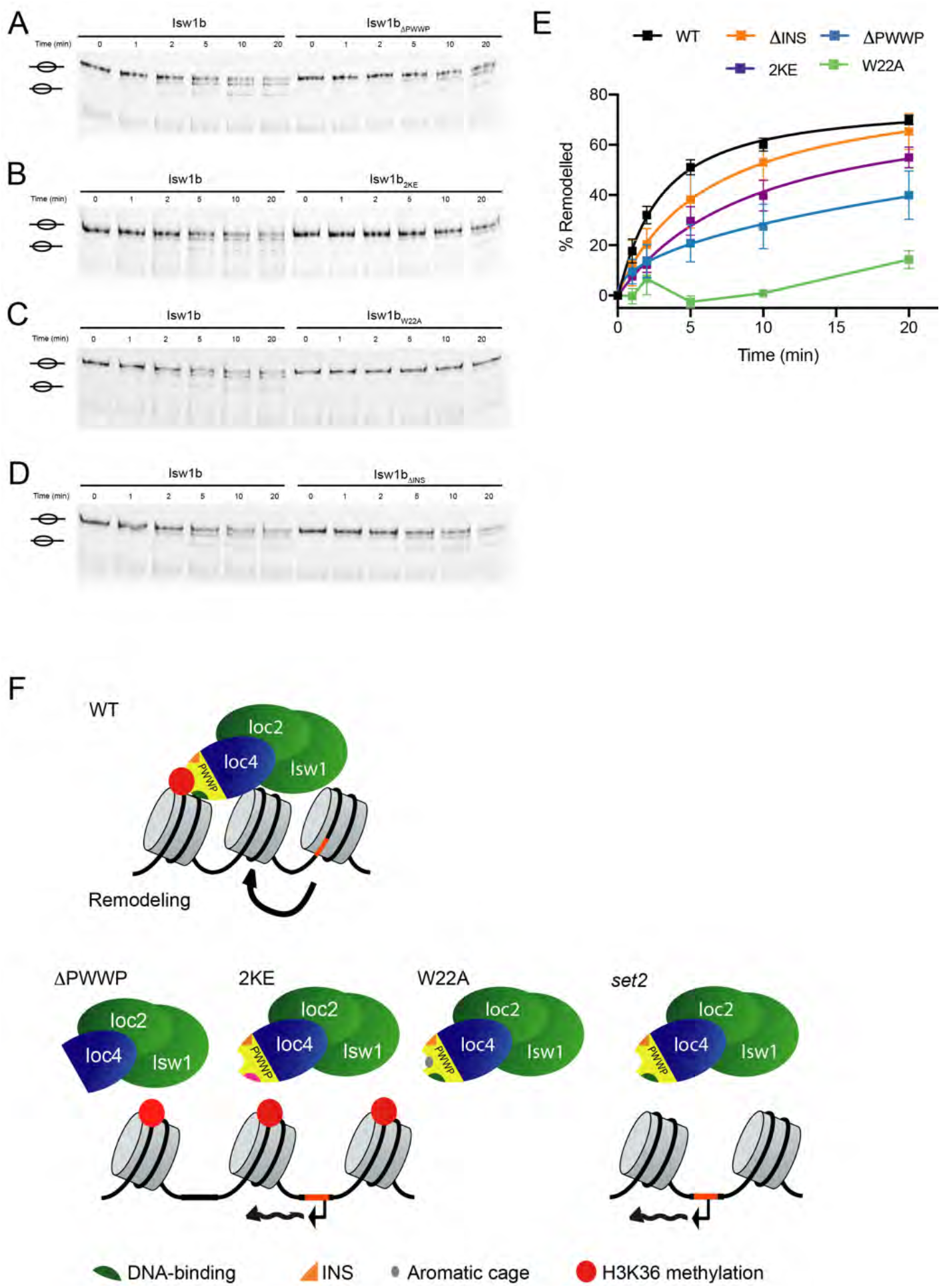
The intact Ioc4 PWWP domain promotes remodelling by Isw1b. **(A-D)** Sliding assays of wildtype and mutant Isw1b remodellers lacking the Ioc4 PWWP domain (A), containing the 2KE (B) or W22A (C) site mutations, or lacking the insertion motif (D). Centrally positioned, MLA-K36Cme3-containing mononucleosomes were remodelled by the addition of remodeller for up to 20 min. Center and side positioned nucleosomes are indicated. **(E)** Quantitation of sliding assays (n=3) shown in (A-D). Bands corresponding to remodelled nucleosomes were quantitated and normalised to their respective input bands (0 min) and plotted as mean ± s.e.m. **(F)** Model for the recruitment of Isw1b through the Ioc4 PWWP domain. In wildtype yeast the Ioc4 PWWP domain simultaneously interacts with H3K36 methylated nucleosomes and the nucleosomal DNA and thereby ensures recruitment of Ioc4 to gene bodies. The remodeller activity of Isw1b maintains proper nucleosome organisation over ORFs which prevents the exposure of promoter-like elements (orange) and the appearance of cryptic transcripts. Without the Ioc4 PWWP domain (ΔPWWP) recruitment of Ioc4 to ORFs is severely curtailed and Isw1b cannot remodel nucleosomes effectively, which disrupts chromatin organisation and results in inappropriate transcription initiation. Interfering with the ability of the PWWP domain to interact with DNA (2KE) has similar effects. The inability of an aromatic cage mutant (W22A) to recognise H3K36me3 or the absence of H3K36 methylation *(set2)* also interferes with Ioc4 recruitment. Interestingly, disruption of PWWP domain binding to methylated H3K36, DNA or both results in the same recruitment defects *in vivo*. The PWWP insertion motif promotes binding of the PWWP domain to DNA, histones and nucleosomes, yet its deletion does not lead to significant changes in Isw1b activity in vitro or in vivo.

While deletion of the insertion motif affects the interactions of the PWWP with DNA, histones and nucleosomes *in vitro*, it had very little effect on the remodelling activity of Ioc4_ΔINS_-containing Isw1b (Fig. 6D,E). These results suggest that the mutant Isw1b_ΔINS_ is able to fulfill remodeller functions *in vivo* in a manner similar to the wildtype, although the insertion motif has a contributory role in remodeller recruitment.

## Discussion

Here, we report the crystal structure of the Ioc4 PWWP domain as well as a detailed analysis of its binding preferences. We have analyzed the PWWP domain on its own as well as in the context of the full-length Ioc4 protein and the entire Isw1b chromatin remodeller, both *in vitro* and *in vivo* (Fig. 6H).

The crystal structure of the Ioc4 PWWP domain exhibits two interesting features that help us understand its biological function. In addition to the preferential binding of the Ioc4 PWWP domain to H3K36 trimethylated nucleosomes via its aromatic cage (Fig. 1,S2), its basic patches are important for interactions with DNA (Fig. 2,3). Mutations within these basic patches lead to a complete loss of DNA and nucleosome binding. Interactions of PWWP domains with DNA have been described also for the PWWP domains of other proteins such as the human LEDGF, DNMT3b and MSH6 proteins as well as Pdp1 and Pdp3 from fission yeast (12,23,24,27,37). Modelling of the Ioc4 PWWP domain onto the cryo-EM structure of LEDGF-PWWP (Fig. 3) supports the notion that different subgroups of PWWP domains interact in similar fashion with nucleosomes by binding across the DNA gyres (27). It also visualises convincingly that K149 and K150 are in close proximity to the nucleosomal DNA and that their mutation leads to a loss of binding of both DNA and nucleosomes.

Interestingly, the Ioc4 PWWP domain also binds to single-stranded RNA and hybrid RNA:DNA molecules with affinities comparable to that of double-stranded DNA (Fig. 2). Whether nucleosomes and ssRNA or RNA:DNA molecules can be bound simultaneously remains unclear. However, it potentially further links Ioc4 and its PWWP domain to elongating RNAPII. In fact, the Isw1b subunits Ioc2 and Isw1 were shown to be able to interact with RNA specifically in RNA immunoprecipitation experiments, while Ioc4 was not investigated (26).

The second interesting feature of the Ioc4 PWWP domain is its remarkably long insertion motif which packs against the β-barrel. Some other PWWP domains also contain short insertions, yet their functions remain unclear. The insertion motif in the Ioc4 PWWP domain contributes both towards one of the basic patches involved in DNA binding as well as an acidic patch promoting PWWP domain binding to histones. Deletion of the insertion motif leads to the destabilization of the interactions between the PWWP domain with DNA, histones and nucleosomes (Fig. 3). Our Ioc4 PWWP-nucleosome model places the insertion motif next to one of the nucleosomal DNA gyres as well as in close proximity to the histone H3 tail, in agreement with our *in vitro* experiments.

In addition to analyzing the PWWP domain on its own we have also evaluated its features in the context of full-length Ioc4 proteins as well as the entire Isw1b remodeller. Comparison with full-length Ioc4 shows that while the PWWP domain is responsible for the preferential binding to K36me3-containing nucleosomes, overall affinities are enhanced ca. 20-fold in the context of full-length Ioc4. Incorporation of Ioc4 into the Isw1b remodeller further increases the affinity of the remodeller for K36 trimethylated nucleosomes. This behaviour agrees with the expectation that the Ioc subunits are responsible for guiding Isw1 remodelers towards target sites.

The integrity of the PWWP domain and its interactions with nucleosomal DNA as well as its association with methylated H3K36 all play important roles for Ioc4 localisation *in vivo*. Disruption of PWWP domain binding to methylated H3K36 in a *set2*Δ mutant, as well as the aromatic cage mutant W22A, unable to recognize H3K36me3, show reduced localization *in vivo*. Similarly, the DNA-binding deficient 2KE mutant exhibits lower recruitment to ORFs. Combining the W22A and 2KE mutations had no further effect on *in vivo* localisation beyond that shown by the single mutants (Fig. 4).

Also, without the Ioc4 PWWP domain Isw1b remodels nucleosomes less effectively (Fig. 6), disrupting chromatin organisation and promoting the production of cryptic and antisense transcripts (Fig. 5). Interfering with the ability of the PWWP domain to interact with DNA (2KE) or bind H3K36me3 (W22A) has similar effects, in agreement with the Ioc4 recruitment data (Fig. 4).

Our previous work has shown that Isw1 and Isw1b in particular are required for the recycling of H3K36me3-modified histones over gene bodies during transcription, thus preventing the incorporation of soluble and highly acetylated histones which predispose yeast cells to the generation of non-coding transcripts. It has been suggested that Isw1b may stabilise existing nucleosomes after RNAPII passage, thereby retaining existing, H3K36 trimethylated histones. The mode of interaction between the Ioc4 PWWP domain and the nucleosome suggested by our model supports this interpretation. Indeed, in agreement with this idea we have some preliminary data to suggest that Ioc4 is able to stabilise nucleosomes *in vitro* under conditions that otherwise lead to their disintegration. Also, nucleosomal arrays in *isw1Δ chd1*Δ cells are known to be severely disrupted (38,39). More recently, a paper from the Clark Lab has shown that the Isw1b remodeller, together with Chd1 is important for resolving closely packed dinucleosomes (10). Failure to do so results in poor nucleosome phasing (10) which in turn could lead to the exposure of cryptic promoters that are usually inaccessible.

A number of PWWP domain-containing proteins have been identified in recent years. They are associated with a variety of protein (complexes) and involved in a diverse set of functions, such as DNA repair (24,40), transcriptional activation (41,42) and repression (3,22,43), DNA methyltransferases (12,44–46), histone lysine methyltransferases (47) and acetyltransferases (48–52) as well as limiting the spreading of heterochromatin (53,54). Many of these PWWP domains preferentially bind methylated histones, with particular emphasis on methylated H3K36 or H4K20 (3,15,43). A positively charged DNA-binding surface has been identified in several other PWWP domains and seems to represent a further common characteristic. The affinities for different PWWP domains and their substrates vary, although all of the domains studied bind nucleosomes with much higher affinities when compared to histone tail peptides. In general, the Ioc4 PWWP domain binds nucleosomes with lower affinity compared to domains from other proteins, such as LEDGF (18,19). However, most binding assays have focused on individual PWWP domains. We show that binding affinities for nucleosomes are greatly enhanced in the context of full-length Ioc4 and the entire Isw1b chromatin remodeller, while the PWWP domain promotes the recognition of K36 methylated nucleosomes and correct targeting for all constructs. Given that PWWP domains are fairly highly conserved, displaying most differences in the length of their insertion motifs and the N-terminal alpha-helical domain, the breadth of diverse molecular functions for these domains may seem somewhat surprising. However, the frequent co-occurrence of PWWP domains with other histone- or DNA-binding domains, such as PHD fingers, bromodomains, AT hooks or chromodomains can account for these differences (3,55) and underlines the importance of studying these domains in context whenever possible.

## Materials & Methods

### Yeast strains and media

All yeast strains used in this study are listed in Table S2. Wildtype *IOC4* was tagged with a 3xFLAG epitope by targeted homologous integration of a PCR product derived from amplification of plasmid p3xFLAG-HIS3 with gene-specific primers. In order to generate yeast strains bearing different mutations of *IOC4*, mutant *IOC4* constructs were first cloned into plasmid p3xFLAG-HIS3, followed by PCR amplification of these cassettes and transformation into wildtype yeast. To generate TAP-tagged yeast strains, the 3xFLAG tag was replaced by targeted homologous integration of a PCR product derived from amplification of plasmid pBS1539 with construct-specific primers. Single deletion of *CHD1* was done by targeted homologous recombination of PCR fragments containing either the hygromycin *(HphB)* or kanamycin *(KanMX)* resistance marker. All strains generated in this study were confirmed by PCR and/or sequencing. Cells were grown at 30°C in YPD (1% yeast extract, 2% bacto-peptone, 2% dextrose) medium.

### Yeast growth assay

Yeast strains were inoculated at an OD_600_ of 0.1 from overnight cultures and grown for ca. 5 hours at 30°C in YPD until they were growing exponentially. Equal numbers of cells were harvested by centrifugation, washed with ddH2O and resuspended at an OD_600_ of 0.5 in ddH_2_O. Five 6-fold dilutions were prepared for each strain and spotted onto YPD plates +/− 1μM propiconazole. Plates were incubated at 30°C for 3-5 days.

### Protein purification

For crystallization the Ioc4 PWWP domain (aa 1-178; Ioc4_PWWP_) was cloned into a modified pMAL-c2X vector (New England Biolabs, Liu *et al*., 2001) in order to produce a fusion protein with an N-terminal maltose-binding protein (MBP). Overexpression of MBP-Ioc4_PWWP_ in *Escherichia coli* BL21 (DE3) was induced by the addition of 0.25 mM IPTG for 20 hours at 16 °C. MBP-Ioc4_PWWP_ was purified to homogeneity using a sequence of amylose resin affinity chromatography, anion exchange and gel filtration chromatography. The PWWP domain of wildtype (aa 1-178) and the PWWP_2KE_ mutant were cloned into a pRSF vector with an N-terminal 6xHis-tag. Both proteins were overexpressed in *E. coli* BL21(DE3) by the addition of 0.25 mM IPTG for 20 hours at 16 °C. Both proteins were purified to homogeneity using nickel-NTA resin, followed by anion exchange chromatography. Full-length Ioc4 (aa1-475), Ioc4_2KE_ and Ioc4_□PWWP_ (aa 179-475) were cloned into a modified pCoofy vector with an N-terminal 6xHis-MBP tag. Proteins were overexpressed in *E. coli* BL21 RIL by addition of 0.25 mM IPTG for 20 hours at 16 °C. Proteins were batch purified using nickel-NTA resin, followed by adsorption chromatography using a heparin column and gel filtration chromatography. Proteins were dialyzed exhaustively against 50 mM phosphate, pH 8.0, 500 mM NaCl, 10% Glycerol, 50 mM Arg, 50 mM Glu, flash-frozen in liquid nitrogen and stored at −80 °C.

TAP-tagged wildtype and mutant Isw1b chromatin remodellers were purified from *Saccharomyces cerevisiae*. Yeast were grown in YPD at 30°C until they reached an OD_600_ of 5-7. Cells were harvested by centrifugation at 5,000 *g* for 15 min and washed twice with cold PBS. Cells were resuspended in 20 ml of TAP Extraction Buffer (40 mM HEPES-KOH, pH7.5, 350 mM NaCl, 10% Glycerol, 0.1% Tween-20, 1 mM PMSF, 2 μg/ml Leupeptin, 1 μg/ml Pepstatin A) and lysed using a freezer mill. Lysates were treated with 100 μl of 10 mg/ml Heparin and 10 μl of Benzonase (25 U/μl, Merck Millipore) for 15 minutes at room temperature before removing cell debris by centrifugation at 31,000 *g* for 20 minutes and ultracentrifugation at 208,000 *g* for 1.5 hours. The supernatant was incubated with pre-washed IgG sepharose (GE Healthcare) and incubated at 4 °C for 3 hours. The resin was washed with TAP extraction buffer for three times 5 min, resuspended with TEV cleavage buffer (10 mM Tris pH 8.0, 10% Glycerol, 150 mM NaCl, 0.1% IGEPAL CA630, 0.5 mM EDTA, 1 mM DTT) and incubated with TEV protease for 16 hours at 4°C. Cleaved protein was collected and applied to calmodulin sepharose (GE Healthcare) pre-washed with binding buffer (10 mM Tris, pH8.0, 150 mM KCl, 1 mM Magnesium acetate, 1 mM Imidazole, 2 mM CaCl2, 10% Glycerol, 0.1% IGEPAL CA630, 1 mM DTT) for 3 hours at 4°C. The resin wash washed with binding buffer for three times 10 min. The purified complexes were eluted with elution buffer (10 mM Tris, pH8.0, 150 mM KCl, 1 mM Magnesium acetate, 1 mM Imidazole, 10 mM EGTA, 10% Glycerol, 0.1% IGEPAL CA630, 0.5 mM DTT), concentrated and subsequently flash frozen in liquid nitrogen and stored at −80°C.

Recombinant *Xenopus laevis* histones were expressed, purified and assembled into core histone octamers as described (56). To generate histone H3 containing either unmethylated or trimethylated K36, tail peptides (aa1-44) containing either K36me0 or K36me3 were ligated onto histone H3_T45C_ and used for subsequent core histone octamer assembly (57). For sliding assays methyl-lysine analogue (MLA) H3_K36C_ nucleosomes chemically modified to resemble either unmethylated or trimethylated H3K36 were used instead, as described previously (8).

### Crystallization and structure determination

Purified MBP-Ioc4_PWWP_ was concentrated to approximately 10 mg/ml and used for crystal screening by sitting-drop vapor diffusion method at 16°C. Needle-shaped crystals grew in reservoir solution containing 0.1 M Potassium thiocyanate, 30% Polyethylene glycol monomethyl ether 2,000. Data were collected on beamline BL17U (SSRF, China, λ = 0.979 Å) and processed using the HKL2000 program suite. Structure of the fusion protein was solved by molecular replace method with the program PHASER, using the MBP structure as a search template. An initial model was automatically built by Phenix Autosol, manually modified with Coot and refined with Phenix Refine. The final model contains two missing fragments, F19 and S43-K91 and has an *R*_work_ of 17.3% and an *Rfree* of 22.9%. Data scaling, refinement, and validation statistics are shown in Table S1.

### GST pull-down assays

GST-tagged Ioc4 constructs were incubated with 2 μg of recombinant, reconstituted histone H3/H4 tetramers and 10 μl of washed glutathione sepharose in buffer B (20 mM Tris-HCl, pH 7.5, 200 mM KCl, 0.1% IGEPAL, 20% Glycerol, 0.2 mM EDTA, 1 mM PMSF, 1 μg/ml pepstatin A, 2 μg/ml leupeptin) for 2.5 hours at 4 °C. Beads were washed with 3x 1 ml of buffer B containing 300 mM KCl, eluted with SDS-PAGE loading buffer and analyzed by SDS-PAGE on 15% gels.

### Reconstitution of recombinant mononucleosomes

DNA fragments measuring either 147 bp or 215 bp and containing the 601 positioning sequence were PCR amplified from pGEM-3Z/601 (58) using Cy5-labeled primers (Table S3). Mononucleosomes were reconstituted from DNA and recombinant *c*ore histone octamers by serial dilution as described (56,59).

### Electromobility shift assays (EMSA)

Binding of Ioc4_PWWP_ to DNA was assessed by EMSA. Double-stranded (ds), Cy5-labeled DNA with a length of 30 bp was prepared by heating and annealing complimentary, single-stranded (ss) DNA (Table S3). Gels were visualised by scanning on a Typhoon FLA9500 Imaging system and quantitated using ImageQuant TL software (GE Healthcare). DNA bands were quantitated for each lane. All lanes containing PWWP were normalised against input lanes containing DNA only. The percentage of DNA bound was expressed as 100 - % DNA for each lane.

EMSA assays with reconstituted, K36me0- or K36me3-containing mononucleosomes were performed as described (56). For binding reactions 15 fmol of reconstituted nucleosomes per reaction were incubated with increasing concentrations (0, 200, 300, 400, 500, 800 nM) of wildtype and mutant PWWP domains. For binding reactions with full-length Ioc4 15 fmol of reconstituted nucleosomes per reaction were incubated with increasing concentrations (0, 6, 12, 18, 24, 35 nM) of wildtype or mutant Ioc4. For binding reactions with unmodified wildtype nucleosomes 30 fmol of reconstituted nucleosomes per reaction were incubated with increasing concentrations (0, 0.2, 1, 2, 4 μM) of wildtype or mutant PWWP. Complexes were separated by electrophoresis on 5.0% native polyacrylamide gels (37.5:1), run in 0.4x TBE, 2% glycerol. Gels were scanned using a Typhoon Imaging FLA9500 system and quantitated using ImageQuant TL software (GE Healthcare). Mononucleosome bands were quantitated for each lane. All lanes containing PWWP were normalised against input lanes containing mononucleosomes only. The percentage of nucleosomes bound was expressed as 100 - % mononucleosomes for each lane.

### Remodelling assays

For nucleosome sliding assays, 10 μl reactions were set up, containing 30 fmol mononucleosomes and 10 fmol Isw1b complex in buffer R (50 mM Tris-HCl, pH8.0, 50 mM KCl, 10 mM MgCl2, 1 mM ATP, 0.1 μg/μl BSA, 5 mM DTT, 0.5 mM PMSF). Reactions were incubated at room temperature and aliquots removed at various time points and stopped by adding 720 ng plasmid DNA, 500 mM KCl. Nucleosomes were resolved by electrophoresis on 7% native polyacrylamide gels (37.5:1) in 0.4x TBE, 2% glycerol. Gels were scanned using a Typhoon FLA9500 Imaging system and quantitated using ImageQuant TL software (GE Healthcare). Bands representing remodelled mononucleosomes were normalised to their respective input lanes.

### Antibodies

The following antibody was used in this study: αFlag M2 (Sigma #F1804).

### Chromatin immunoprecipitation assays

For ChIPs of FLAG-tagged Ioc4 yeast strains were grown in 200 ml of YPD at 30 °C, crosslinked and processed for ChIP as described earlier (60,61). FLAG-tagged Ioc4 was immunoprecipitated and processed as described before (60,61). Three biological repeats were done for all experiments and used for subsequent ChIP-chip analysis.

### ChIP-qPCR analysis

Immunoprecipitated DNA was quantitated by qPCR using the PowerTrack SYBR Green Master Mix (Thermo Fisher Scientific) and a QuantStudio 5 Real-Time PCR System (Applied Biosystems). Primers are listed in Table S3. The mean signal was calculated for each experiment and normalised against input samples at each primer positions as internal controls. Input-normalised values were further corrected for variation by normalising against the mean signals for two control regions *(STE3*, subtelomeric region on chromosome V *(ChrV)*.

### ChIP-chip microarray analysis

ChIP-chip assays for the genome-wide distribution of FLAG-tagged Ioc4 were performed as described previously (8), using 8×60K yeast genome DNA arrays (Agilent, Array #031697) with an average probe spacing of ca. 200 bp. 20-50 ng of input and IP samples were used for double T7 linear amplification and labelling. Inputs were labelled with Cy3 dye and IPs with Cy5 dye. 4 μg of input and IP were combined and used for hybridisation. Arrays were scanned (Agilent DNA Microarray Scanner Model G2505B) and extracted using Feature Extraction software (Agilent) and normalised using median normalisation in R software.

### Data analysis

The normalised data were analyzed using a modified gene average analysis (8). ORFs were subdivided into 14 equal sized bins each. Intergenic regions (480 bp up- and downstream of genes) were allocated into three bins each. Microarray enrichment ratios [log_2_(IP/Input)] for each probe were assigned to the closest bin. For whole-genome average gene plots all probes within a bin were averaged and plotted as mean ± standard error (SEM). Genes not regulated by RNAPII, including tRNA and snRNA genes as well as the majority of dubious ORFs (ca. 450 genes) were removed from the analysis.

### Isolation of total RNA

Yeast strains were grown in YPD at 30°C until they reached an OD_600_ of 0.8. Total RNA was isolated using acid phenol extraction as described previously (62). RNA quality and quantity were assessed by UV spectroscopy using a NanoDrop 2000 (Thermo Scientific).

### Northern blots

Northern blotting and hybridization were done as described previously (63). 20 μg of total RNA were used to assess the cryptic transcript phenotype. Blots were exposed onto phosphorimaging screens and scanned using a Typhoon FLA9500 Imaging system.

### Strand-specific multiplex RT-qPCR

Total RNA was treated with TURBO DNA-free DNase I (ThermoScientific) to remove genomic DNA according to the manufacturer’s instructions. 3.5 μg of DNase-treated RNA were used for subsequent reverse transcription (RT) reactions containing 2 pmol of each gene-specific primer, 60 U of SuperScript III (ThermoScientific) and 0.3 μg of Actinomycin D (AppliChem) to prevent antisense cDNA artefacts. Two RT reactions containing “forward” primers (Table S3) annealing to antisense transcripts derived from target genes *VTH2* and *YEN1*, or *FAA2* and *ARO80* were set up. Each reaction also contained 2 pmol “reverse” primer annealing to the canonical reference *ACT1* transcript. Annealing was done at 70°C for 10 minutes. First strand synthesis was performed at 55°C for one hour, followed by heat inactivation at 70°C for 15 min. All samples were quantitated by qPCR using the PowerTrack SYBR Green Master Mix (Thermo Fisher Scientific) and a QuantStudio 5 Real-Time PCR System (Applied Biosystems). Ct-values were first normalised to those of *ACT1*. Subsequently, results for mutant yeast strains were normalized to the mean signal for wildtype yeast samples using the comparative CT (ΔΔCT) method. Five biological replicates were performed for each strain.

## Accession codes

Gene Expression Omnibus: ChIP-chip data sets have been deposited with accession number GSE165984. Crystal coordinates were deposited with the Protein Data Bank (accession code 7E29).

## Acknowledgements

We thank Julia Schluckebier for excellent technical assistance. Experiments were done using equipment in the BioPhysics Core Facility of the Biomedical Center, LMU. Crystallography data were collected at beamline BL17U at the SSRF, China. H.L. was supported by science and technology planning project of Shenzhen (JCYJ20180307155005435). M.S. was supported by the Deutsche Forschungsgmeinschaft (SFB1064, SM426/1-1) and the Friedrich-Bauer-Stiftung (07/15). P.V. was supported by the Wellcome Trust [104175/Z/14/Z] and through funding from the European Research Council (ERC-STG grant 639253). The Wellcome Centre for Cell Biology was supported by core funding from the Wellcome Trust [203149]. We are grateful to the Edinburgh Protein Production Facility (EPPF) for their support. The EPPF was supported by the Wellcome Trust through a Multi-User Equipment grant [101527/Z/13/Z].

## Contributions

J.L. purified and crystallised the Ioc4-PWWP domain. H.L. determined and analyzed the structure. L.B. purified all other Ioc4 and Isw1b remodeller constructs and performed all *in vitro* experiments. K.W. and P.V. provided peptide-ligated histone octamers. A.R.A. performed RT-qPCR experiments. M.S. performed all other *in vivo* experiments. M.S. and M.G. performed bioinformatics analysis. M.S., H.L. and Y.L. supervised research. M.S. wrote the manuscript with input from all authors.

## Supporting Information

**Table S1.** Statistics of crystallographic analysis. *Values in parentheses are for highest-resolution shell.

|  |  |
| --- | --- |
| **Data collection** |  |
| Space group | P63 |
| Cell dimension |  |
| a,b,c(Å) | 153.303, 153.303, 42.004 |
| $\alpha,\beta,\gamma(^{\circ})$ | 90.000, 90.000, 120.000, |
| Resolution (Å) | 50.0-2.30(2.34 -2.30) * |
| $R_{\text{meq}}$ (%) | 10.9/(70.6) |
| $I/\sigma I$ | 35.5(5.6) |
| Completeness (%) | 99.9(100.0) |
| Redundancy | 7.9(7.8) |
| **Refinement** |  |
| Resolution (Å) | 2.3 |
| Total No. reflection / free | 25341/1995 |
| $R_{\text{work}} / R_{\text{free}}$ | 0.173/0.229 |
| r.m.s.d. bonds/angles (Å) | 0.008/1.117 |
| Protein/solvent atoms | 3910/ 300 |
| Average B-factors(Å<sup>2</sup>) | 35.451 |
| **Ramachandran plot statistics** |  |
| Most favorable | 98.57% |
| Additionally allowed | 1.22% |
| Disallowed | 0.20% |

**Table S2.** Yeast strains used in this study.

| Strain | Parental | Genotype | Source |
| --- | --- | --- | --- |
| BY4741 | BY4741 | *MATa his3Δ1 leu2Δ0 met15Δ0 ura3Δ0* | Open Biosystems |
| YMS061 | BY4741 | *MATa his3Δ1 leu2Δ0 met15Δ0 ura3Δ0 IOC4-TAP::HIS3* | Open Biosystems |
| YMS087 | BY4741 | *MATa his3Δ1 leu2Δ0 met15Δ0 ura3Δ0 IOC4-3xFlag::loxP set2Δ::LEU2* | M. Smolle (1) |
| YMS205 | BY4741 | *MATa his3Δ1 leu2Δ0 met15Δ0 ura3Δ0 ioc4Δ::HIS3 chd1Δ::KanMX* | This study |
| YMS254 | BY4741 | *MATa his3Δ1 leu2Δ0 met15Δ0 ura3Δ0 IOC4-3xFlag::HIS3* | This study |
| YMS255 | BY4741 | *MATa his3Δ1 leu2Δ0 met15Δ0 ura3Δ0 IOC4<sub>Δ43-105</sub>-3xFlag::HIS3* | This study |
| YMS262 | BY4741 | *MATa his3Δ1 leu2Δ0 met15Δ0 ura3Δ0 IOC4-3xFlag::HIS3 chd1Δ::HphB* | This study |
| YMS263 | BY4741 | *MATa his3Δ1 leu2Δ0 met15Δ0 ura3Δ0 IOC4<sub>Δ43-105</sub>-3xFlag::HIS3 chd1Δ::HphB* | This study |
| YMS264 | BY4741 | *MATa his3Δ1 leu2Δ0 met15Δ0 ura3Δ0 IOC4<sub>Δ1-178</sub>-3xFlag::HIS3* | This study |
| YMS265 | BY4741 | *MATa his3Δ1 leu2Δ0 met15Δ0 ura3Δ0 IOC4<sub>K149E K150E</sub>-3xFlag::HIS3* | This study |
| YMS266 | BY4741 | *MATa his3Δ1 leu2Δ0 met15Δ0 ura3Δ0 IOC4<sub>Δ1-178</sub>-3xFlag::HIS3 chd1Δ::HphB* | This study |
| YMS267 | BY4741 | *MATa his3Δ1 leu2Δ0 met15Δ0 ura3Δ0 IOC4<sub>K149E K150E</sub>-3xFlag::HIS3 chd1Δ::HphB* | This study |
| YMS333 | BY4741 | *MATa his3Δ1 leu2Δ0 met15Δ0 ura3Δ0 IOC4<sub>Δ43-105</sub> -TAP::URA3* | This study |
| YMS334 | BY4741 | *MATa his3Δ1 leu2Δ0 met15Δ0 ura3Δ0 IOC4<sub>Δ1-178</sub> -TAP::URA3* | This study |
| YMS335 | BY4741 | *MATa his3Δ1 leu2Δ0 met15Δ0 ura3Δ0 IOC4<sub>K149E K150E</sub> -TAP::URA3* | This study |
| YMS376 | BY4741 | *MATa his3Δ1 leu2Δ0 met15Δ0 ura3Δ0 IOC4<sub>W22A</sub>-3xFlag::HIS3* | This study |
| YMS378 | BY4741 | *MATa his3Δ1 leu2Δ0 met15Δ0 ura3Δ0 IOC4<sub>W22A</sub>-TAP::URA3* | This study |
| YMS379 | BY4741 | *MATa his3Δ1 leu2Δ0 met15Δ0 ura3Δ0 IOC4<sub>W22A K149E K150E</sub>-3xFlag::HIS3* | This study |

**Table S3.**
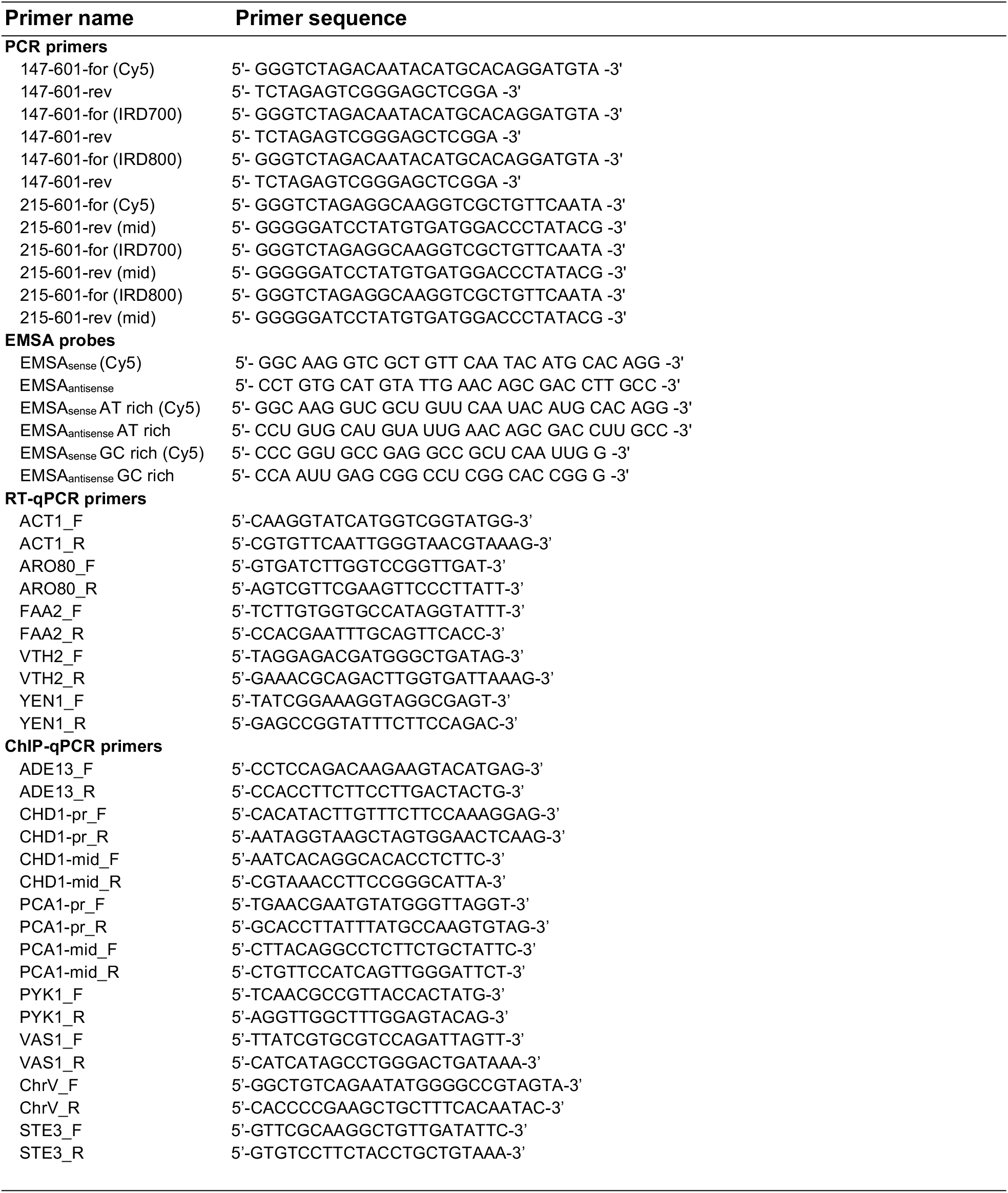
Primers used in this study.

**Figure S1.**
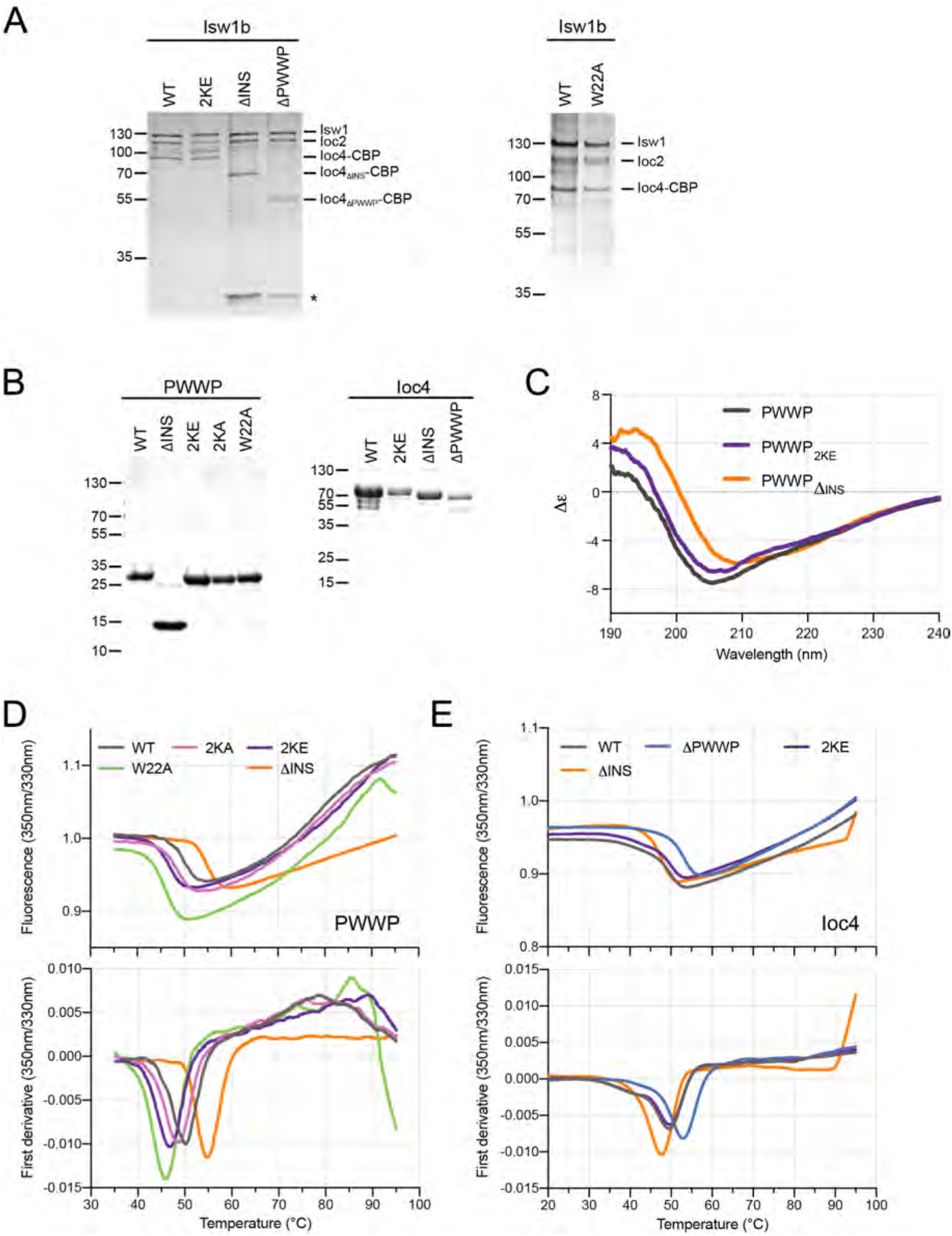
Protein purification and stability. **(A)** Silver stained SDS-PAGE of TAP-purified wildtype and mutant Isw1b remodeler complexes. TEV protease was identified by mass spectrometry and is indicated (*). **(B)** SDS-PAGE analysis of purified, wildtype and mutant PWWP and Ioc4 constructs. Proteins were stained with Coomassie Blue. **(C)** Circular dichroism analysis of pufified PWWP constructs shows that all proteins are folded. **(D,E)** Nano differential scanning fluorimetry (DSF) results for purified PWWP **(D)** and Ioc4 **(E)** proteins. The PWWP constructs were analysed using the Nanotemper Tycho, the Ioc4 proteins were run on the Nanotemper Prometheus. Results are not directly comparable due to different settings for the temperature ramp.

**Figure S2.**
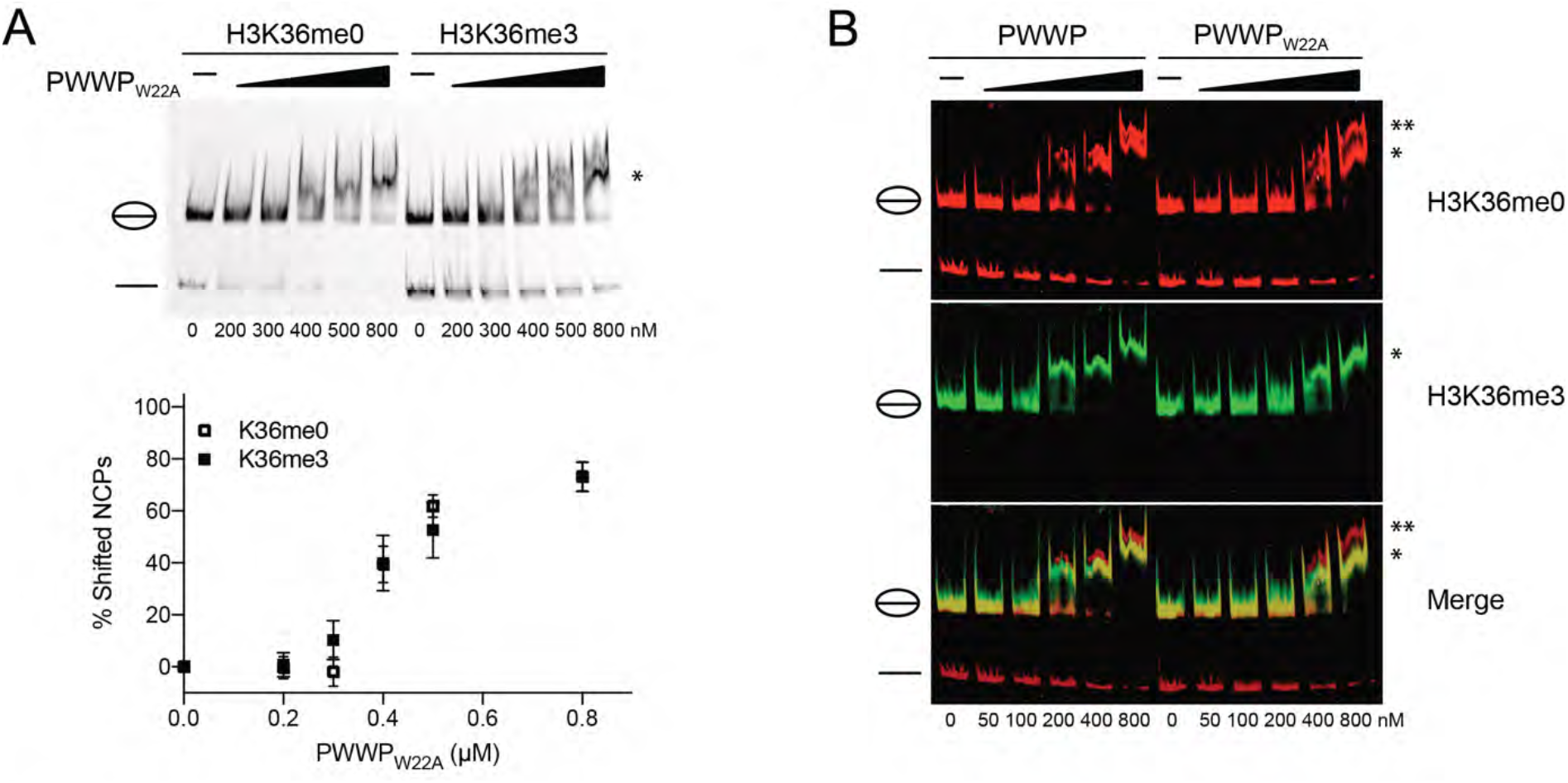
The PWWP aromatic cage is important for H3K36me3 recognition. EMSAs **(A)** and competitive EMSAs **(B)** were performed to assess the ability of the aromatic cage mutant W22A to specifically recognise H3K36 trimethylated nucleosomes. As expected, mutation of W22 led to a loss of discrimination in both settings. Furthermore, the mutant PWWP domain displayed lower affinities towards nucleosomes in general, when compared to the wildtype domain. The positions of the NCP and free DNA bands on the gels are indicated, as are the bands denoting the complexes formed by the PWWP proteins with NCPs (*) as well as with free DNA (**).

**Figure S3.**
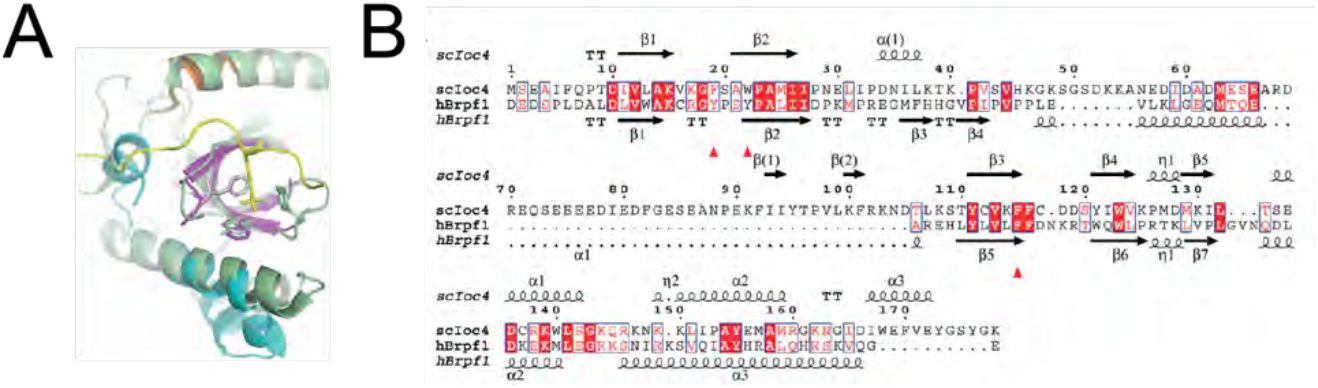
Structural and sequence conservation of PWWP domains **(A)** Overlay of the PWWP domains from Ioc4 (colored as in Fig. 1H) and hBrpf1 (light green). The H3K36me3-containing peptide is shown in yellow. Residues forming the aromatic cage are shown as sticks. **(B)** Sequence alignment of PWWP domains from Ioc4 (scIoc4) and hBrpf1. The alignment was generated with Clustal Omega and displayed with ESPript (http://espript.ibcp.fr/). Secondary structure elements of the Ioc4 and Brpf1 PWWP domains are marked at the top and bottom of the alignment, respectively. Identical residues are highlighted in red. Aromatic cage residues accommodating the histone trimethyllysine are labeled with red triangles.

**Figure S4.**
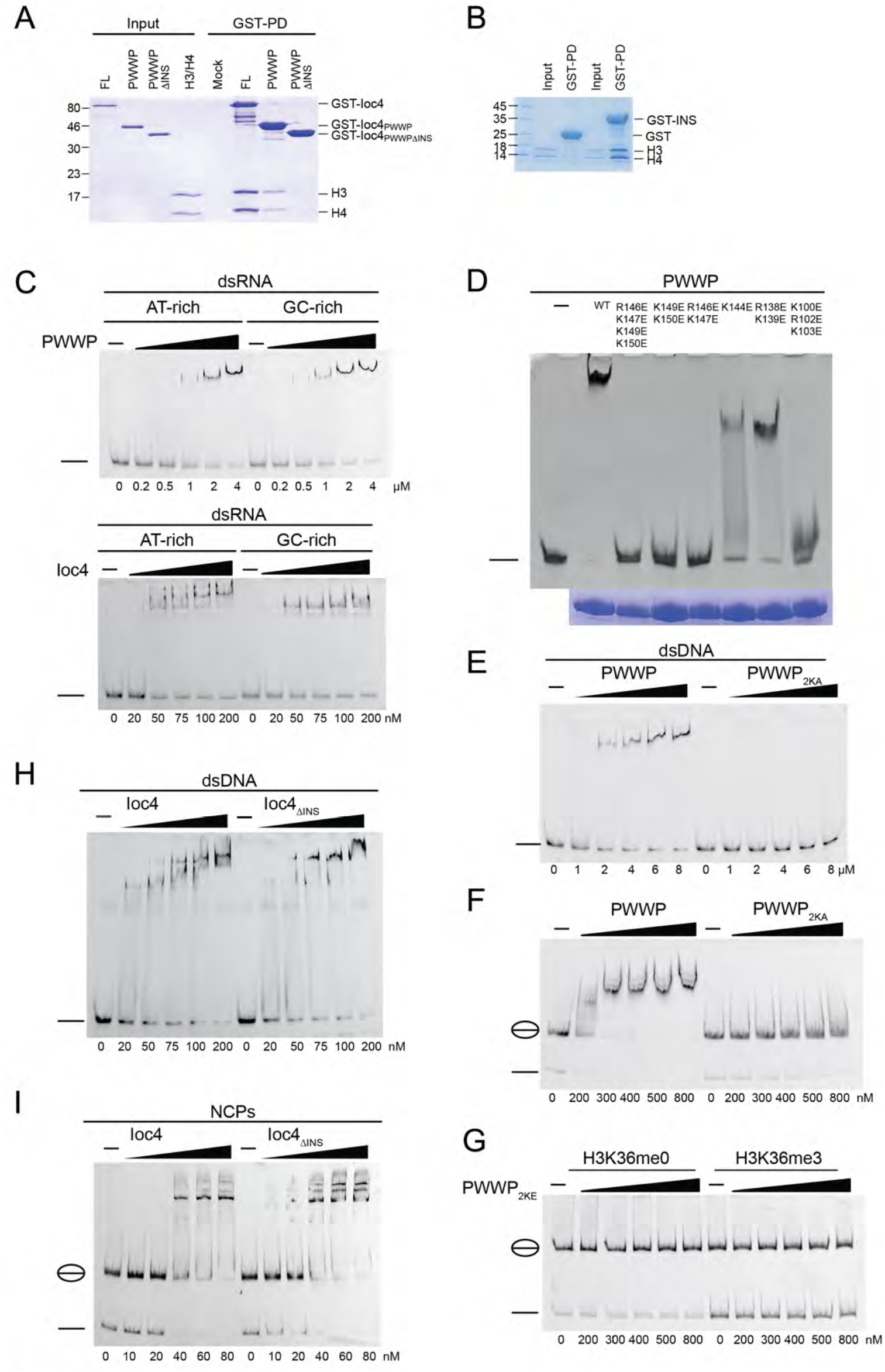
Interaction of Ioc4 and Ioc4_PWWP_ constructs with histones and nucleosomes **(A)** Pull-down assays of GST-tagged Ioc4, PWWP and PWWP without its insertion motif (PWWP_ΔINS_) with histone H3/H4 tetramers. **(B)** Pull-down assays of GST-tagged insertion motif (INS) with histone H3/H4 tetramers. **(C)** EMSA of wildtype PWWP and Ioc4 with dsRNA. **(D)** Wildtype and mutant PWWP domains were purified, used to set up binding reactions with double-stranded DNA (30 bp) and analyzed by EMSA. The DNA bands are indicated by a line. Equal protein loading is shown. **(E)** EMSA of wildtype PWWP and PWWP_2KA_ with dsDNA (30 bp). **(F)** EMSA of wildtype PWWP and PWWP_2KA_ with unmodified NCPs. **(G)** EMSA of PWWP_2KE_ with unmethylated and trimethylated H3K36 NCPs. **(H)** EMSA of Ioc4 and Ioc4_ΔINS_ with dsDNA. **(I)** EMSA of Ioc4 and Ioc4_ΔINS_ with NCPs. Free NCPs and/or DNA are indicated.

**Figure S5.**
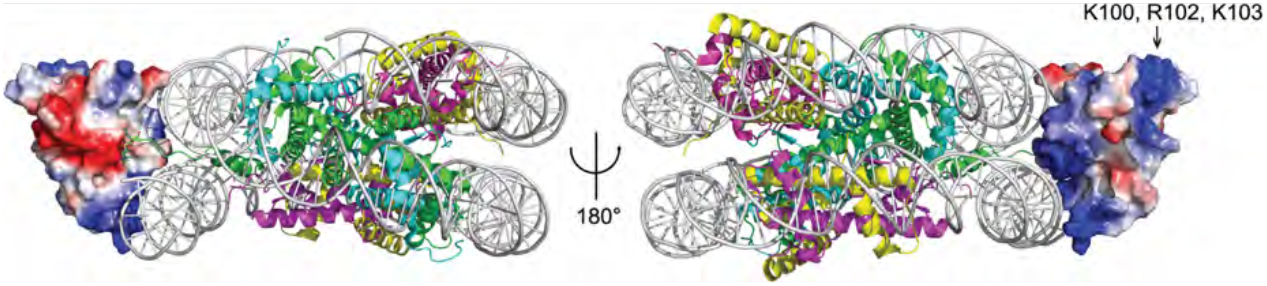
Model of Ioc4 PWWP binding to NCP. Homology Model based on Cramer et al. (2). The electrostatic surface of the Ioc4-PWWP is shown. Basic residues (K100, R102, K103) part of the insertion motif are indicated.

**Figure S6.**
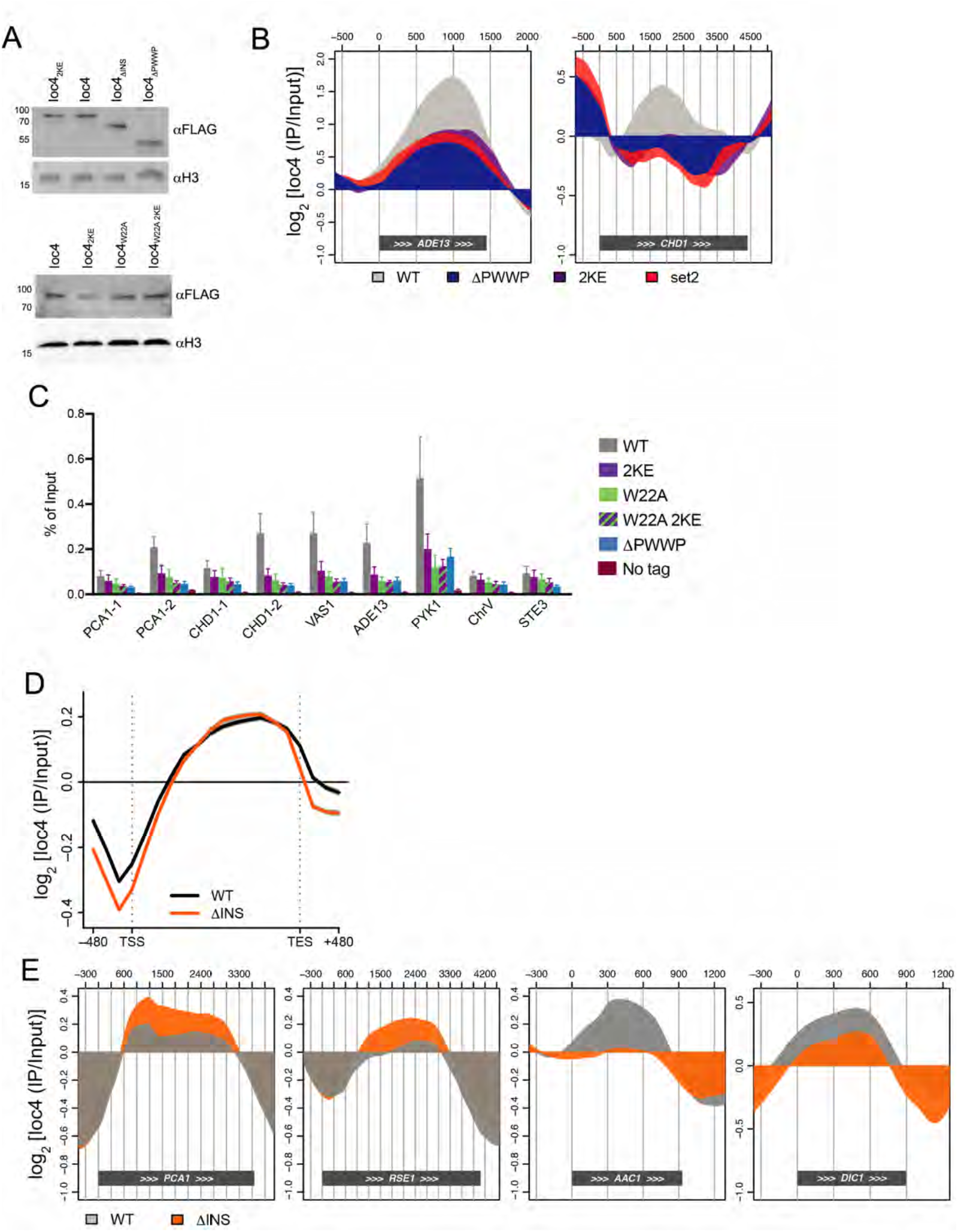
Ioc4 recruitment to chromatin. **(A)** Western blot of whole cell extracts from yeast strains expressing wildtype or mutant 3xFlag tagged Ioc4. Histone H3 was used as a loading control. **(B)** Localisation of wildtype and mutant Ioc4 over individual genes. (**C**) ChIP-qPCR experiments were performed for wildtype and mutant IOC4-3xFlag strains. BY4741 was used as an untagged control. Localisation was determined over promoter and ORF regions. *STE3* and *ChrV* served as control regions. All ChIP signals were normalised to input. **(D)** Metagene analysis of ChIP-chip experiments using yeast genome tiling arrays. Whole-genome average data (n=6451 genes) for three independent experiment were plotted as mean ± s.e.m. (gray) for wildtype Ioc4 and Ioc4_ΔINS_. (**E)** Localisation of wildtype Ioc4 and Ioc4_ΔINS_ over individual genes.

## References

1. Hughes, A.L. and Rando, O.J. (2014) Mechanisms underlying nucleosome positioning in vivo. Annual review of biophysics, 43, 41–63.

2. Smolle, M. and Workman, J.L. (2013) Transcription-associated histone modifications and cryptic transcription. Biochim Biophys Acta, 1829, 84–97.

3. Qin, S. and Min, J. (2014) Structure and function of the nucleosome-binding PWWP domain. Trends Biochem Sci, 39, 536–547.

4. Narlikar, G.J., Sundaramoorthy, R. and Owen-Hughes, T. (2013) Mechanisms and Functions of ATP-Dependent Chromatin-Remodeling Enzymes. Cell, 154, 490–503.

5. Elfring, L.K., Deuring, R., McCallum, C.M., Peterson, C.L. and Tamkun, J.W. (1994) Identification and characterization of Drosophila relatives of the yeast transcriptional activator SNF2/SWI2. Mol Cell Biol, 14, 2225–2234.

6. Tsukiyama, T., Palmer, J., Landel, C.C., Shiloach, J. and Wu, C. (1999) Characterization of the imitation switch subfamily of ATP-dependent chromatin-remodeling factors in Saccharomyces cerevisiae. Genes Dev, 13, 686–697.

7. Vary, J.C., Jr., Gangaraju, V.K., Qin, J., Landel, C.C., Kooperberg, C., Bartholomew, B. and Tsukiyama, T. (2003) Yeast Isw1p forms two separable complexes in vivo. Mol Cell Biol, 23, 80–91.

8. Smolle, M., Venkatesh, S., Gogol, M.M., Li, H., Zhang, Y., Florens, L., Washburn, M.P. and Workman, J.L. (2012) Chromatin remodelers Isw1 and Chd1 maintain chromatin structure during transcription by preventing histone exchange. Nat Struct Mol Biol, 19, 884–892.

9. Venkatesh, S., Smolle, M., Li, H., Gogol, M., Saint, M., Kumar, S., Natarajan, K. and Workman, J.L. (2012) Set2 methylation of histone H3 lysine36 suppresses histone exchange on transcribed genes. Nature, 489, 452–455.

10. Eriksson, P.R. and Clark, D.J. (2021) The yeast ISW1b ATP-dependent chromatin remodeler is critical for nucleosome spacing and dinucleosome resolution. Sci Rep, 11, 4195.

11. Maurer-Stroh, S., Dickens, N.J., Hughes-Davies, L., Kouzarides, T., Eisenhaber, F. and Ponting, C.P. (2003) The Tudor domain ‘Royal Family’: Tudor, plant Agenet, Chromo, PWWP and MBT domains. ‘Trends Biochem Sci, 28, 69–74.

12. Qiu, C., Sawada, K., Zhang, X. and Cheng, X. (2002) The PWWP domain of mammalian DNA methyltransferase Dnmt3b defines a new family of DNA-binding folds. Nat Struct Biol, 9, 217–224.

13. Ge, Y.Z., Pu, M.T., Gowher, H., Wu, H.P., Ding, J.P., Jeltsch, A. and Xu, G.L. (2004) Chromatin targeting of de novo DNA methyltransferases by the PWWP domain. J Biol Chem, 279, 25447–25454.

14. Wu, H., Zeng, H., Lam, R., Tempel, W., Amaya, M.F., Xu, C., Dombrovski, L., Qiu, W., Wang, Y. and Min, J. (2011) Structural and histone binding ability characterizations of human PWWP domains. PLoS One, 6, e18919.

15. Wang, Y., Reddy, B., Thompson, J., Wang, H., Noma, K., Yates, J.R., 3rd and Jia, S. (2009) Regulation of Set9-mediated H4K20 methylation by a PWWP domain protein. Mol Cell, 33, 428–437.

16. Maltby, V.E., Martin, B.J., Schulze, J.M., Johnson, I., Hentrich, T., Sharma, A., Kobor, M.S. and Howe, L. (2012) Histone H3 lysine 36 methylation targets the Isw1b remodeling complex to chromatin. Mol Cell Biol, 32, 3479–3485.

17. Pinskaya, M., Nair, A., Clynes, D., Morillon, A. and Mellor, J. (2009) Nucleosome remodeling and transcriptional repression are distinct functions of Isw1 in Saccharomyces cerevisiae. Mol Cell Biol, 29, 2419–2430.

18. van Nuland, R., van Schaik, F.M., Simonis, M., van Heesch, S., Cuppen, E., Boelens, R., Timmers, H.M. and van Ingen, H. (2013) Nucleosomal DNA binding drives the recognition of H3K36-methylated nucleosomes by the PSIP1-PWWP domain. Epigenetics & chromatin, 6, 12.

19. Eidahl, J.O., Crowe, B.L., North, J.A., McKee, C.J., Shkriabai, N., Feng, L., Plumb, M., Graham, R.L., Gorelick, R.J., Hess, S. et al. (2013) Structural basis for high-affinity binding of LEDGF PWWP to mononucleosomes. Nucleic Acids Res.

20. Tian, W., Yan, P., Xu, N., Chakravorty, A., Liefke, R., Xi, Q. and Wang, Z. (2019) The HRP3 PWWP domain recognizes the minor groove of double-stranded DNA and recruits HRP3 to chromatin. Nucleic Acids Res, 47, 5436–5448.

21. Rona, G.B., Eleutherio, E.C.A. and Pinheiro, A.S. (2016) PWWP domains and their modes of sensing DNA and histone methylated lysines. Biophys Rev, 8, 63–74.

22. Yang, J. and Everett, A.D. (2007) Hepatoma-derived growth factor binds DNA through the N-terminal PWWP domain. BMC Mol Biol, 8, 101.

23. Qiu, Y., Zhang, W., Zhao, C., Wang, Y., Wang, W., Zhang, J., Zhang, Z., Li, G., Shi, Y., Tu, X. et al. (2012) Solution structure of the Pdp1 PWWP domain reveals its unique binding sites for methylated H4K20 and DNA. Biochem J, 442, 527–538.

24. Laguri, C., Duband-Goulet, I., Friedrich, N., Axt, M., Belin, P., Callebaut, I., Gilquin, B., Zinn-Justin, S. and Couprie, J. (2008) Human mismatch repair protein MSH6 contains a PWWP domain that targets double stranded DNA. Biochemistry, 47, 6199–6207.

25. Lukasik, S.M., Cierpicki, T., Borloz, M., Grembecka, J., Everett, A. and Bushweller, J.H. (2006) High resolution structure of the HDGF PWWP domain: a potential DNA binding domain. Protein Sci, 15, 314–323.

26. Babour, A., Shen, Q., Dos-Santos, J., Murray, S., Gay, A., Challal, D., Fasken, M., Palancade, B., Corbett, A., Libri, D. et al. (2016) The Chromatin Remodeler ISW1 Is a Quality Control Factor that Surveys Nuclear mRNP Biogenesis. Cell, 167, 1201–1214 e1215.

27. Wang, H., Farnung, L., Dienemann, C. and Cramer, P. (2020) Structure of H3K36-methylated nucleosome-PWWP complex reveals multivalent cross-gyre binding. Nat Struct Mol Biol, 27, 8–13.

28. Chabbert, C.D., Adjalley, S.H., Klaus, B., Fritsch, E.S., Gupta, I., Pelechano, V. and Steinmetz, L.M. (2015) A high-throughput ChIP-Seq for large-scale chromatin studies. Molecular Systems Biology, 11, 777.

29. Sayou, C., Millán-Zambrano, G., Santos-Rosa, H., Petfalski, E., Robson, S., Houseley, J., Kouzarides, T. and Tollervey, D. (2017) RNA Binding by Histone Methyltransferases Set1 and Set2. Molecular and Cellular Biology, 37, e00165–00117.

30. Cole, H.A., Ocampo, J., Iben, J.R., Chereji, R.V. and Clark, D.J. (2014) Heavy transcription of yeast genes correlates with differential loss of histone H2B relative to H4 and queued RNA polymerases. Nucleic Acids Res, 42, 12512–12522.

31. Weiner, A., Hughes, A., Yassour, M., Rando, O.J. and Friedman, N. (2010) High-resolution nucleosome mapping reveals transcription-dependent promoter packaging. Genome Res, 20, 90–100.

32. Kireeva, M.L., Walter, W., Tchernajenko, V., Bondarenko, V., Kashlev, M. and Studitsky, V.M. (2002) Nucleosome remodeling induced by RNA polymerase II: loss of the H2A/H2B dimer during transcription. Mol Cell, 9, 541–552.

33. Belotserkovskaya, R., Oh, S., Bondarenko, V.A., Orphanides, G., Studitsky, V.M. and Reinberg, D. (2003) FACT facilitates transcription-dependent nucleosome alteration. Science, 301, 1090–1093.

34. Guan, M., Xia, P., Tian, M., Chen, D. and Zhang, X. (2020) Molecular fingerprints of conazoles via functional genomic profiling of Saccharomyces cerevisiae. Toxicol In Vitro, 69, 104998.

35. Quan, T.K. and Hartzog, G.A. (2010) Histone H3K4 and K36 methylation, Chd1 and Rpd3S oppose the functions of Saccharomyces cerevisiae Spt4-Spt5 in transcription. Genetics, 184, 321–334.

36. Stockdale, C., Flaus, A., Ferreira, H. and Owen-Hughes, T. (2006) Analysis of nucleosome repositioning by yeast ISWI and Chd1 chromatin remodeling complexes. J Biol Chem, 281, 16279–16288.

37. Georgescu, P.R., Capella, M., Fischer-Burkart, S. and Braun, S. (2020) The euchromatic histone mark H3K36me3 preserves heterochromatin through sequestration of an acetyltransferase complex in fission yeast. Microb Cell, 7, 80–92.

38. Gkikopoulos, T., Schofield, P., Singh, V., Pinskaya, M., Mellor, J., Smolle, M., Workman, J.L., Barton, G.J. and Owen-Hughes, T. (2011) A role for Snf2-related nucleosome-spacing enzymes in genome-wide nucleosome organization. Science Now York, N.Y, 333, 1758–1760.

39. Ocampo, J., Chereji, R.V., Eriksson, P.R. and Clark, D.J. (2016) The ISW1 and CHD1 ATP-dependent chromatin remodelers compete to set nucleosome spacing in vivo. Nucleic Acids Res, 44, 4625–4635.

40. Huang, Y., Gu, L. and Li, G.M. (2018) H3K36me3-mediated mismatch repair preferentially protects actively transcribed genes from mutation. J Biol Chem, 293, 7811–7823.

41. Zhu, X., Lan, B., Yi, X., He, C., Dang, L., Zhou, X., Lu, Y., Sun, Y., Liu, Z., Bai, X. et al. (2020) HRP2-DPF3a-BAF complex coordinates histone modification and chromatin remodeling to regulate myogenic gene transcription. Nucleic Acids Res, 48, 6563–6582.

42. Fei, J., Ishii, H., Hoeksema, M.A., Meitinger, F., Kassavetis, G.A., Glass, C.K., Ren, B. and Kadonaga, J.T. (2018) NDF, a nucleosome-destabilizing factor that facilitates transcription through nucleosomes. Genes Dev, 32, 682–694.

43. Wen, H., Li, Y., Xi, Y., Jiang, S., Stratton, S., Peng, D., Tanaka, K., Ren, Y., Xia, Z., Wu, J. et al. (2014) ZMYND11 links histone H3.3K36me3 to transcription elongation and tumour suppression. Nature, 508, 263–268.

44. Chen, T., Tsujimoto, N. and Li, E. (2004) The PWWP domain of Dnmt3a and Dnmt3b is required for directing DNA methylation to the major satellite repeats at pericentric heterochromatin. Mol Cell Biol, 24, 9048–9058.

45. Otani, J., Nankumo, T., Arita, K., Inamoto, S., Ariyoshi, M. and Shirakawa, M. (2009) Structural basis for recognition of H3K4 methylation status by the DNA methyltransferase 3A ATRX-DNMT3-DNMT3L domain. EMBO Rep, 10, 1235–1241.

46. Weinberg, D.N., Papillon-Cavanagh, S., Chen, H., Yue, Y., Chen, X., Rajagopalan, K.N., Horth, C., McGuire, J.T., Xu, X., Nikbakht, H. et al. (2019) The histone mark H3K36me2 recruits DNMT3A and shapes the intergenic DNA methylation landscape. Nature, 573, 281–286.

47. He, C., Li, F., Zhang, J., Wu, J. and Shi, Y. (2013) The methyltransferase NSD3 has chromatin-binding motifs, PHD5-C5HCH, that are distinct from other NSD (nuclear receptor SET domain) family members in their histone H3 recognition. J Biol Chem, 288, 4692–4703.

48. Vezzoli, A., Bonadies, N., Allen, M.D., Freund, S.M., Santiveri, C.M., Kvinlaug, B.T., Huntly, B.J., Gottgens, B. and Bycroft, M. (2010) Molecular basis of histone H3K36me3 recognition by the PWWP domain of Brpf1. Nat Struct Mol Biol, 17, 617–619.

49. Laue, K., Daujat, S., Crump, J.G., Plaster, N., Roehl, H.H., Tubingen Screen, C., Kimmel, C.B., Schneider, R. and Hammerschmidt, M. (2008) The multidomain protein Brpf1 binds histones and is required for Hox gene expression and segmental identity. Development, 135, 1935–1946.

50. Liu, L., Qin, S., Zhang, J., Ji, P., Shi, Y. and Wu, J. (2012) Solution structure of an atypical PHD finger in BRPF2 and its interaction with DNA. J Struct Biol, 180, 165–173.

51. Poplawski, A., Hu, K., Lee, W., Natesan, S., Peng, D., Carlson, S., Shi, X., Balaz, S., Markley, J.L. and Glass, K.C. (2014) Molecular insights into the recognition of N-terminal histone modifications by the BRPF1 bromodomain. J Mol Biol, 426, 1661–1676.

52. Qin, S., Jin, L., Zhang, J., Liu, L., Ji, P., Wu, M., Wu, J. and Shi, Y. (2011) Recognition of unmodified histone H3 by the first PHD finger of bromodomain-PHD finger protein 2 provides insights into the regulation of histone acetyltransferases monocytic leukemic zinc-finger protein (MOZ) and MOZ-related factor (MORF). J Biol Chem, 286, 36944–36955.

53. Dou, K., Liu, Y., Zhang, Y., Wang, C., Huang, Y. and Zhang, Z.Z. (2020) Drosophila P75 safeguards oogenesis by preventing H3K9me2 spreading. J Genet Genomics, 47, 187–199.

54. Albig, C., Wang, C., Dann, G.P., Wojcik, F., Schauer, T., Krause, S., Maenner, S., Cai, W., Li, Y., Girton, J. et al. (2019) JASPer controls interphase histone H3S10 phosphorylation by chromosomal kinase JIL-1 in Drosophila. Nat Commun, 10, 5343.

55. Savitsky, P., Krojer, T., Fujisawa, T., Lambert, J.P., Picaud, S., Wang, C.Y., Shanle, E.K., Krajewski, K., Friedrichsen, H., Kanapin, A. et al. (2016) Multivalent Histone and DNA Engagement by a PHD/BRD/PWWP Triple Reader Cassette Recruits ZMYND8 to K14ac-Rich Chromatin. Cell Rep, 17, 2724–2737.

56. Li, B., Gogol, M., Carey, M., Lee, D., Seidel, C. and Workman, J.L. (2007) Combined action of PHD and chromo domains directs the Rpd3S HDAC to transcribed chromatin. Science (New York, N.Y, 316, 1050–1054.

57. Voigt, P., LeRoy, G., Drury, W.J., 3rd, Zee, B.M., Son, J., Beck, D.B., Young, N.L., Garcia, B.A. and Reinberg, D. (2012) Asymmetrically modified nucleosomes. Cell, 151, 181–193.

58. Thastrom, A., Bingham, L.M. and Widom, J. (2004) Nucleosomal locations of dominant DNA sequence motifs for histone-DNA interactions and nucleosome positioning. J Mol Biol, 338, 695–709.

59. Li, B., Jackson, J., Simon, M.D., Fleharty, B., Gogol, M., Seidel, C., Workman, J.L. and Shilatifard, A. (2009) Histone H3 lysine 36 di-methylation (H3K36ME2) is sufficient to recruit the Rpd3S histone deacetylase complex and to repress spurious transcription. J Biol Chem, 284, 7970–7976.

60. Strahl-Bolsinger, S., Hecht, A., Luo, K. and Grunstein, M. (1997) SIR2 and SIR4 interactions differ in core and extended telomeric heterochromatin in yeast. Genes Dev, 11, 83–93.

61. Li, B. and Reese, J.C. (2001) Ssn6-Tup1 regulates RNR3 by positioning nucleosomes and affecting the chromatin structure at the upstream repression sequence. J Biol Chem, 276, 33788–33797.

62. Li, B., Gogol, M., Carey, M., Pattenden, S.G., Seidel, C. and Workman, J.L. (2007) Infrequently transcribed long genes depend on the Set2/Rpd3S pathway for accurate transcription. Genes Dev, 21, 1422–1430.

63. Carrozza, M.J., Li, B., Florens, L., Suganuma, T., Swanson, S.K., Lee, K.K., Shia, W.J., Anderson, S., Yates, J., Washburn, M.P. et al. (2005) Histone H3 methylation by Set2 directs deacetylation of coding regions by Rpd3S to suppress spurious intragenic transcription. Cell, 123, 581–592.

